# A feather star is born: embryonic development and nervous system organization in the crinoid *Antedon mediterranea*

**DOI:** 10.1101/2024.01.30.577964

**Authors:** S. Mercurio, G. Gattoni, G. Scarì, M. Ascagni, B. Barzaghi, M. R. Elphick, E. Benito-Gutiérrez, R. Pennati

## Abstract

**Background:** Crinoids belong to the phylum Echinodermata, marine invertebrates with a highly derived pentaradial body plan. As the only living members of the Pelmatozoa, the sister group to other extant echinoderms, crinoids are in a key phylogenetic position to reconstruct the evolutionary history of this phylum. However, the development of crinoids has been scarcely investigated, limiting their potential for comparative studies. Many crinoids are difficult to collect in the wild and embryo manipulation is challenging. Conversely, the Mediterranean feather star *Antedon mediterranea* can be found in shallow waters and has been used for experimental studies, most notably to investigate regeneration.

**Results:** The aim here was to establish *A. mediterranea* as an experimental system for developmental biology. To accomplish this, we set up a method for culturing embryos *in vitro* from zygote to hatching larva stage that allowed us to define a developmental timeline and a standardized staging system for this species. We then optimized protocols to characterize the development of the main structures of the feather star body plan, using a combination of microscopy techniques and whole mount immunohistochemistry and *in situ* hybridization chain reaction. Focusing on the nervous system, we show that the larval apical organ includes a combination of serotonergic, GABAergic and glutamatergic neurons that form under the influence of a conserved anterior molecular signature. The larval neural plexus is instead composed of glutamatergic neurons and develops during the formation of the ciliary bands. Larval neurons disappear at metamorphosis, and the ectoneural and entoneural components of the adult nervous system develop early in post-metamorphic stages. Furthermore, the oral ectoderm that contains the ectoneural system acquires an “anterior” signature expressing *Six3/6* and *Lhx2/9* orthologs.

**Conclusions:** Our results deepen our knowledge on crinoid development and provide new techniques to investigate feather star embryogenesis, promoting the use of *A. mediterranea* in developmental and evolutionary biology. This in turn will pave the way for the inclusion of crinoids in comparative studies to understand the origin of the echinoderm body plan and clarify many unanswered questions on deuterostome evolution.

## Background

Echinoderms are some of the most enigmatic animals among deuterostomes (the taxon that includes echinoderms, hemichordates and chordates). They display a highly-derived adult body plan, characterized by pentaradial symmetry, a calcitic endoskeleton, a coelom-derived water vascular system and a peculiar nervous system, in which the presence of a centralized integrative center is under debate [1, 2]. Modern echinoderms are divided into two main taxa: the Eleuterozoa, comprising four extant classes (asteroids, ophiuroids, echinoids and holothuroids), and the Pelmatozoa, of which crinoids are the only living members [3]. In turn, crinoids can be sub-divided into stalked sea lilies, which remain sessile after metamorphosis, and stalkless feather stars that detach from their stalk after post-larval development and become free-living adults [4, 5].

As the sister group to the rest of the echinoderms, crinoids are in a key phylogenetic position to reconstruct the evolutionary history of the phylum [6]. However, knowledge of crinoid biology and development remains scarce compared to the wealth of information on the life cycle of eleutherozoans. Sea urchins are traditional models in developmental biology, and their early embryogenesis and larval development have been thoroughly investigated [7, 8]. With the advent of molecular biology, eleutherozoans have been used to study gene regulatory networks and their role in the specification of cell types [9–11]. More recently, optimized culturing protocols for research and commercial purposes have made it possible to follow their development through metamorphosis and study juvenile stages [12–14]. Conversely, crinoid embryogenesis and growth have been scarcely investigated. Apart from early morphological descriptions [15, 16], only few studies on the development of these animals have been reported to date [17–20], despite the increasing interest in crinoid biology within the scientific community [21, 4, 22, 23].

Crinoids have swimming ciliated lecithotrophic larvae: sea lilies have an early dipleurula-like stage followed by a barrel-shaped doliolaria, while in feather stars only a doliolaria stage is observed [24, 18, 17]. It was shown previously that doliolaria larvae have a nervous system characterized by a diffuse basiepithelial neural plexus and an anterior serotonergic apical organ, located below a tufted apical pit [4, 19]. Apical organs are a conserved feature of echinoderm development and have been identified in all eleutherozoan taxa. The molecular networks leading to the differentiation of larval neural structures have been dissected in many eleutherozoan models [7, 11, 25]; conversely, the development and cell type composition of the crinoid nervous system have not been studied in detail. After a short larval phase, the doliolaria settles on the seabed and undergoes a gradual metamorphosis, during which the larval tissues are rearranged to form a transient cystidean stage that develops into a sessile pentacrinoid [17, 26, 4]. The majority of recent studies on crinoid development have focused on post-embryonic stages [22, 23, 27], while the molecular cues guiding embryogenesis still remain mostly unexplored. This is partly due to difficulties in collecting wild specimens and in manipulating crinoid gametes and embryos. Crinoids are more abundant in the Indo-Pacific region, where many stalkless crinoids inhabit coastal shallow waters. They are also common in cold waters at depths between 15 and 150 m, while sea lilies are generally regarded as deep sea animals [16, 28]. Moreover, crinoid reproductive seasons are extremely variable, being mostly population-specific, and cues controlling spawning are largely unknown [4, 17, 29, 30].

*Antedon mediterranea* is a Mediterranean feather star traditionally used for regeneration studies [31, 31, 32]. It is an external brooding animal [33], and embryos are retained on females’ genital pinnules until they hatch. Embryogenesis occurs in tight association with the parental individual, making early embryos difficult to access and experimentally manipulate, consequently hindering the use of *Antedon* in developmental biology and evo-devo research [4, 17]. To overcome this obstacle and establish *A. mediterranea* as a new model system, here we set up a method for culturing embryos *in vitro* from zygote to hatching larva stage and established a standardized staging system of feather star development based on the main features that we described for each developmental phase. We then optimized whole-mount immunohistochemistry and *in situ* hybridization chain reaction (HCR) protocols to follow the origin and development of the main larval structures. Finally, we focused on the nervous system and characterized cell type populations and conserved molecular signals in larval and post-metamorphic stages. By providing the first analysis of gene co-expression in crinoids, together with the localization of neurotransmitters, we identified a complex apical organ in the feather star larva that develops within a highly conserved anterior domain. Taken together, our results deepen knowledge of crinoid development and provide a set of techniques that will facilitate use of the crinoid *A. mediterranea* as an experimental model in developmental and evolutionary studies.

## Results

### Spawning induction, embryo collection and embryo culture *in vitro*

We found that the most efficient way to induce *A. mediterranea* spawning in aquaria involves exposing ripe animals to environmental stressors, and in particular to a strong light source (by means of a torch). During the reproductive season, we observed that male individuals are particularly sensitive to light, and promptly respond to light cues by releasing sperm in the water. Female spawning generally occurred after sperm was released by males. Gamete spawning and fertilization occurred within a few minutes after stimulation. While embryos normally remain attached to the females’ genital pinnules, we found that it is possible to detach hundreds of zygotes from females by transferring the into small glass crystallizers (100 ml) filled with seawater (SW) for about half an hour. This appeared to be a stressful condition for females, which responded by releasing most of the zygotes. Embryos were then collected from the bottom of the crystallizers and transferred to plastic petri dishes containing filtered sea water (see Materials and Methods for details), where they were successfully reared (Fig. 1). Frequent water changes and prompt removal of dead specimens were key to ensure survival and good health of the embryos, which under these conditions reached the hatched larva stage in about 100 hpf (hours post fertilization) (17±1°C; Fig. 1 J-L) at a hatching rate of 92.5 ± 1.8%. To ensure that the isolation of the embryos from the mother did not have an impact on their development, we jointly monitored the development of embryos raised in petri-dishes and those attached to adults’ genital pinnules (Fig. 1 A, D, G). We found no developmental differences between embryos growing *in vitro* in petri dishes and those developing attached to the genital pinnules, which reached the hatching stage at the same time (Fig. 1 A-I).

**Fig. 1.**
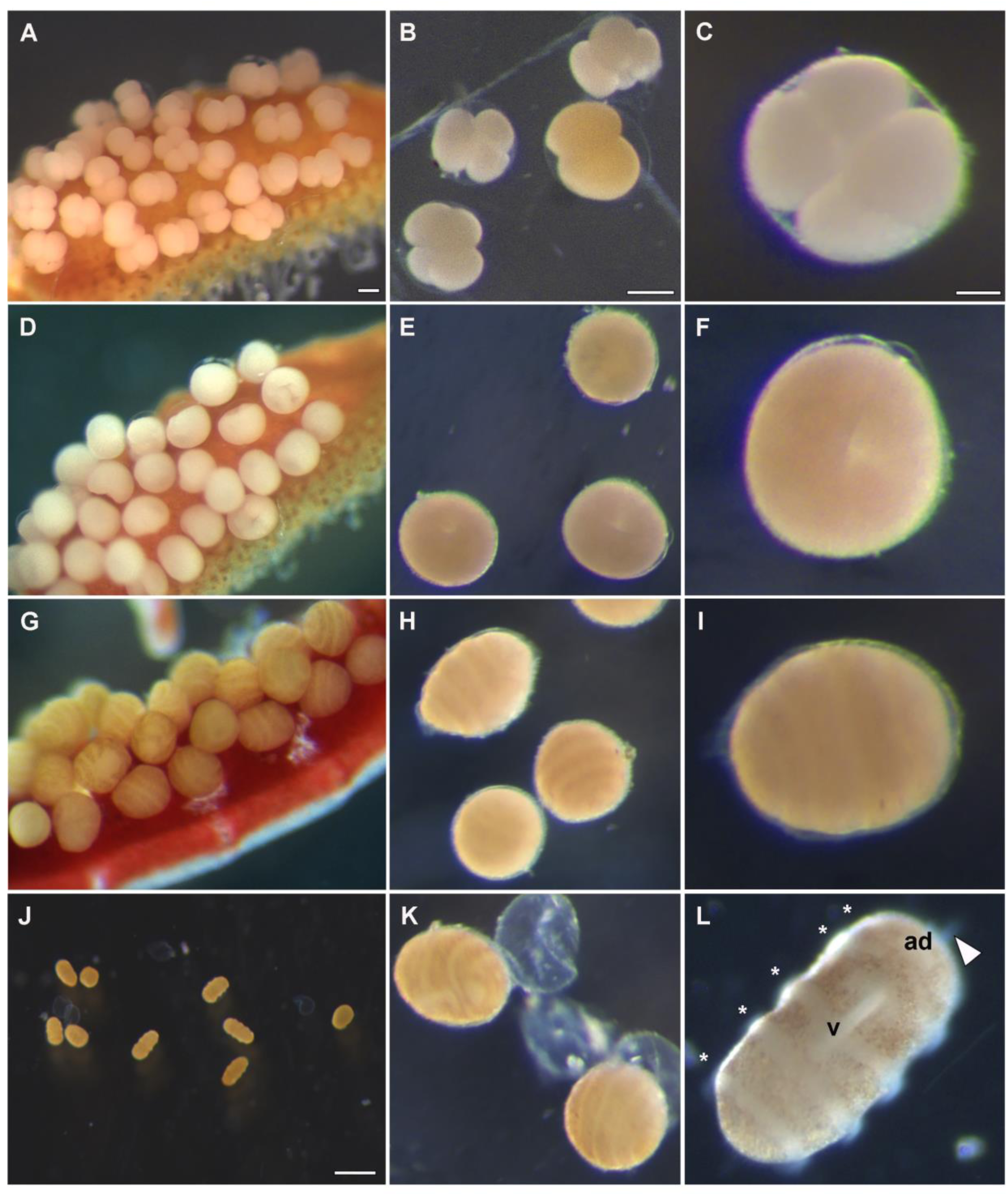
*A. mediterranea* embryogenesis on parental pinnules and *in vitro*. Embryos attached to adults’ genital pinnules in the aquaria (A, D and G) develop synchronously with those maintained *in vitro* (B, C, E, F, H, I, J, L). **A-C**) 4-cell stage; **D-F**) gastrula stage (24 hpf); **G-I**) pre-hatching larvae (72 hpf); **J-L)** hatching (**K**; 100 hpf)) and swimming doliolaria larvae (**J** and **L**) *in vitro*. Legend: * = ciliary band; arrowhead = apical tuft; ad = adhesive pit; v = vestibulum. Scale bars: A = 100 µm; B = 100 µm; C) 50 µm and J = 500 µm.

### Timing and main features of *A. mediterranea* developmental stages

The embryogenesis of *A. mediterranea* was previously described in classical microscopy studies of the late nineteenth and early twentieth century, often accompanied by beautiful drawings of selected developmental stages [15, 16, 34–36]. Here, we expand on these early works by characterizing all the main embryonic events up to larval hatching using modern microscopy techniques, and by providing a standardized staging system of *A. mediterranea* embryogenesis based on easily identifiable developmental features. In parallel, we characterize the pattern of cell divisions in post-cleavage stages, using a newly-optimized protocol of whole mount immunofluorescence to label mitotic nuclei using an antibody specific to phosphorylated histone 3 (PhH3) (Suppl. Fig. 1) [37]. Overall, our standardized staging system spans through 12 pre-metamorphic developmental stages including: zygote stage (fertilization – 2 hpf), 6 cleavage stages (2 – 9 hpf), gastrulation stage (9 – 36 hpf), uniformly ciliated stage (36 – 48 hpf), band formation stage (48 – 72 hpf), pre-hatching doliolaria (72 – 100 hpf), swimming doliolaria (≥100 hpf) (Fig. 2).

**Fig. 2.**
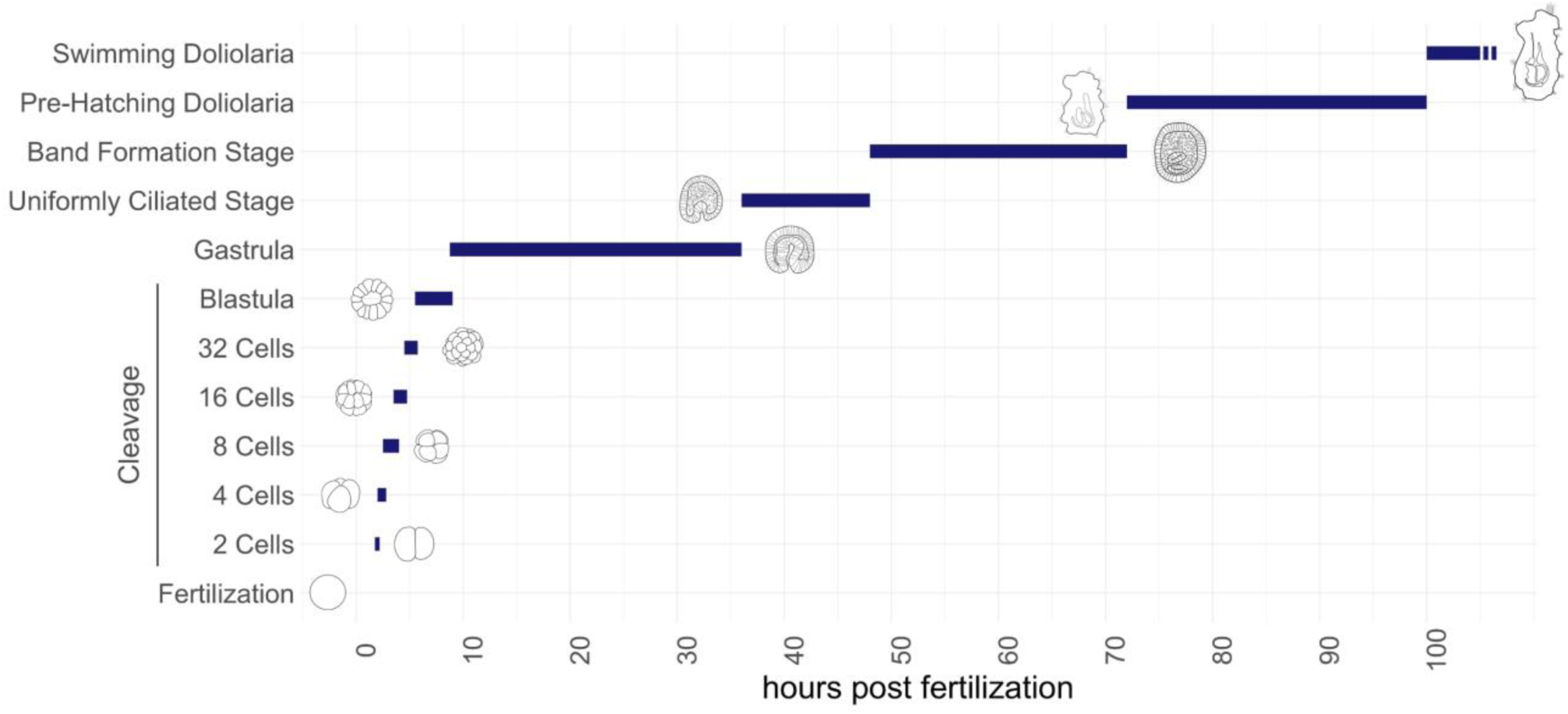
Developmental timeline of *A. mediterranea* embryogenesis. Graphical representation of *A. mediterranea* development at 17±1°C: zygote (fertilization – 2 hpf), 2-cell stage (2 – 2.5 hpf), 4-cell stage (2.5 – 3.5 hpf), 8-cell stage (3.5 – 4 hpf), 16-cell stage (4 – 5 hpf), 32-cell stage (5 – 6 hpf), blastula stage (6 – 9 hpf), gastrulation (9 – 36 hpf), uniformly ciliated stage (36 – 48 hpf), band formation stage (48 – 72 hpf), pre-hatching doliolaria (72 – 100 hpf), swimming doliolaria (≥100 hpf).

*Zygote* (0 – 2 hpf at 17±1°C). The mesolecithical egg was spherical, about 220 µm in diameter (Fig. 3 A). Upon fertilization, an ornamented membrane covered the embryo until hatching. This membrane appeared spiny by light microscopy, but SEM analysis revealed that the structure was folded in thin ridges irregularly distributed along the surface (Suppl. Fig. 2 A-D).

**Fig. 3.**
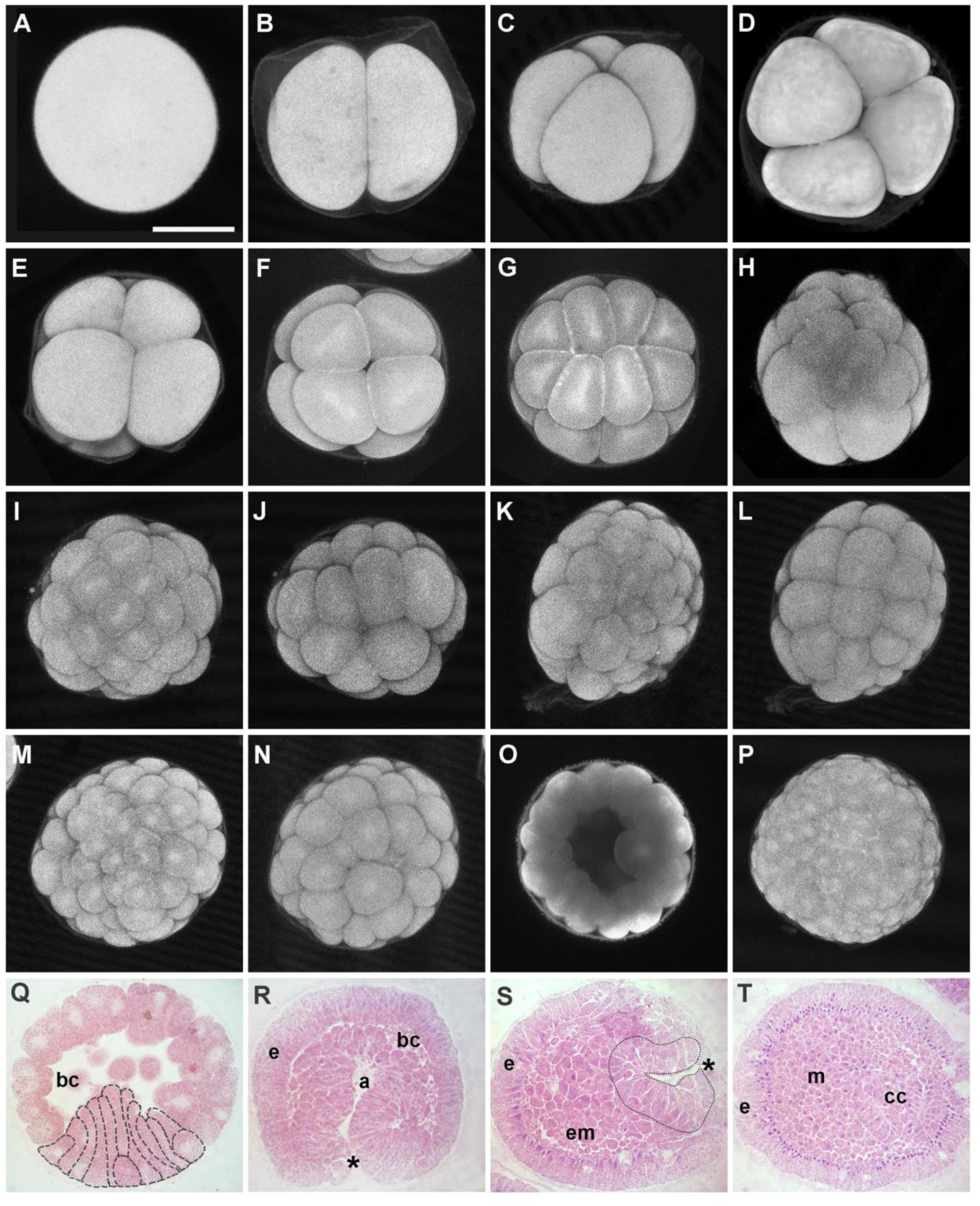
Morphological characterization of *A. mediterranea* embryogenesis. **A-P**) Confocal z-projections of embryos labelled with phalloidin in the cleavage period: **A**) zygote (1 hpf); **B**) 2-cell stage (2 hpf); **C**) 4-cell stage (2.5 hpf), lateral view; **D**) 4-cell stage, frontal view; **E**) 8-cell stage (3.5 hpf), lateral view, animal pole on the top; **F**) 8-cell stage, animal view; **G**) 16-cell stage (4 hpf), animal view; 24-cell stage, lateral view; **I** and **J**) the two sides of a 32-cell stage (5 hpf); **K-L**) the two sides of a 48-cell stage; **M-N**) the two sides of a 64-cell stage (6 hpf); **O**) mid-sagittal projection optical sections of a blastula stage (6 hpf), showing the internal cavity; **P**) initial gastrula (9 hpf). **Q-T**) Light microscopy of *A. mediterranea* embryos undergoing gastrulation: **Q**) sagittal section of an initial gastrula (9 hpf) in which elongating cells at the vegetative pole (outlined with dashed lines) are observable; **R**) sagittal section of a mid-gastrula (20 hpf), with asterisk indicating the blastopore; **S**) late-gastrula (36 hpf), the blastopore is closed and the archenteron is narrow (outlined with dotted line); **T**) ciliated larva (48 hpf) in which coelomic cavities (cc) are developing. Legend: a = archenteron; ab: animal blastomeres; bc = blastocoel; cc = coelomic cavities; e = ectoderm; em = entomesoderm; m = mesoderm; vb: vegetal blastomeres. Scale bar = 100 µm.

*Cleavage period* (∼2 – 9 hpf at 17±1°C). The first cell division occurred at ∼2 hpf and split the zygote into two nearly equal blastomeres (Fig. 3 B). The second cleavage plane ran meridian to the first, and resulted in four cells of similar size (Fig. 3 C, D; ∼2.5 hpf). The third mitotic division separated animal and vegetal poles. The cleavage plane was parallel to the equator but closer to animal pole, thus diving the embryos in four smaller (animal) and four larger (vegetal) blastomeres (Fig. 3 E, F; ∼3.5 hpf). The next divisions occurred first at the animal pole and then in the vegetal one. Most embryos reached the 16-cell stage 4 hpf and consisted of eight smaller animal cells and eight larger vegetal cells (Fig. 3 G). The following cleavage plane ran equatorially and divided the eight animal cells into 16 animal blastomeres of almost equal size, leading to the 24-cell stage, composed of eight large vegetal cells on which two layers of eight small cells were positioned (Fig. 3 H). About 5 hpf, the division of the vegetal blastomeres was completed and the embryo reached the 32-cell stage (Fig. 3 I, J). Rearrangements of cell positions occurred broadly, resulting in embryos with slightly different appearances. Inside the embryo, a wide blastocoel started to form. The following division at the animal pole resulted in an almost spherical embryo of 48 cells (Fig. 3 K, L) which soon reached the 64-cell stage (Fig. 3 M, N; ∼6 hpf), corresponding to the beginning of the blastula period (Fig. 3 O, P, Suppl. Fig. 2). During all the cleavage phase, cell divisions were generally characterized by a marked asynchrony between animal and vegetal pole, with cells dividing faster on the animal side. Moreover, a moderate level of developmental asynchrony was observed among embryos of the same batch, in which the same division did not happen exactly at the same time. This asynchrony among embryos became even more evident from the blastula stage onwards (Fig. 2).

*Gastrulation period* (∼9 hpf – 36 hpf at 17±1°C). At the beginning of this stage, the blastomeres at the vegetal pole of the embryo changed their morphology: the most vegetal columnar cells seemed to undergo apical constriction, elongating toward the blastocoel cavity and acquiring a flask shape (Fig. 3 Q). About 10 hpf, invagination of the ento-mesodermal layer began at the vegetal side and at 20 hpf embryos reached a mid-gastrula stage. At this stage, the ento-mesoderm extended through the blastocoel, almost reaching the ectodermal layer at the animal pole. At the same time, small mesenchymal cells detached from the ento-mesodermal layer and started filling the blastocoel cavity. The ectodermal cells retained their columnar shape, while near the blastopore the ento-mesodermal blastomeres showed a characteristic flask shape (Fig. 3 R). During the next few hours, mesenchyme cells continued to fill the embryo cavity and at ∼36 hpf the blastopore closed in most of the samples and the archenteron assumed the shape of a narrow sac (Fig. 3 S). Through gastrulation, cells in active proliferation were distributed all over the embryos (Suppl. Fig. 1 A).

*Uniformly ciliated stage* (∼36 – 48 hpf at 17±1°C). The formation of mesenchyme continued, and the blastocoel was no longer distinguishable. At the posterior end of the embryo, the archenteron divided first into two vesicles, the entero-hydrocoel and the somatocoel, and then the hydrocoel and the axocoel further separated from the enteric sac (Fig. 3 T, Fig. 4 F, Fig. 5 B, C). Together with the development of the coelomic cavities, the ectodermal cells enveloped the entire embryo and completed their transition into a uniformly ciliated epithelium (Fig. 4 B). At this stage, immunoreactivity to the proliferation marker PhH3 was more concentrated in ectodermal cells (Suppl. Fig. 1 B). The ectoderm of these embryos consisted of columnar cells regularly arranged around the mesodermal tissue (Fig. 3 T). Most of these cells presented basal nuclei in resting stage, while mitotic signal was consistently detected in cells with nuclei localized in the central or apical region of the cell (Suppl. Fig. 1 B).

**Fig. 4.**
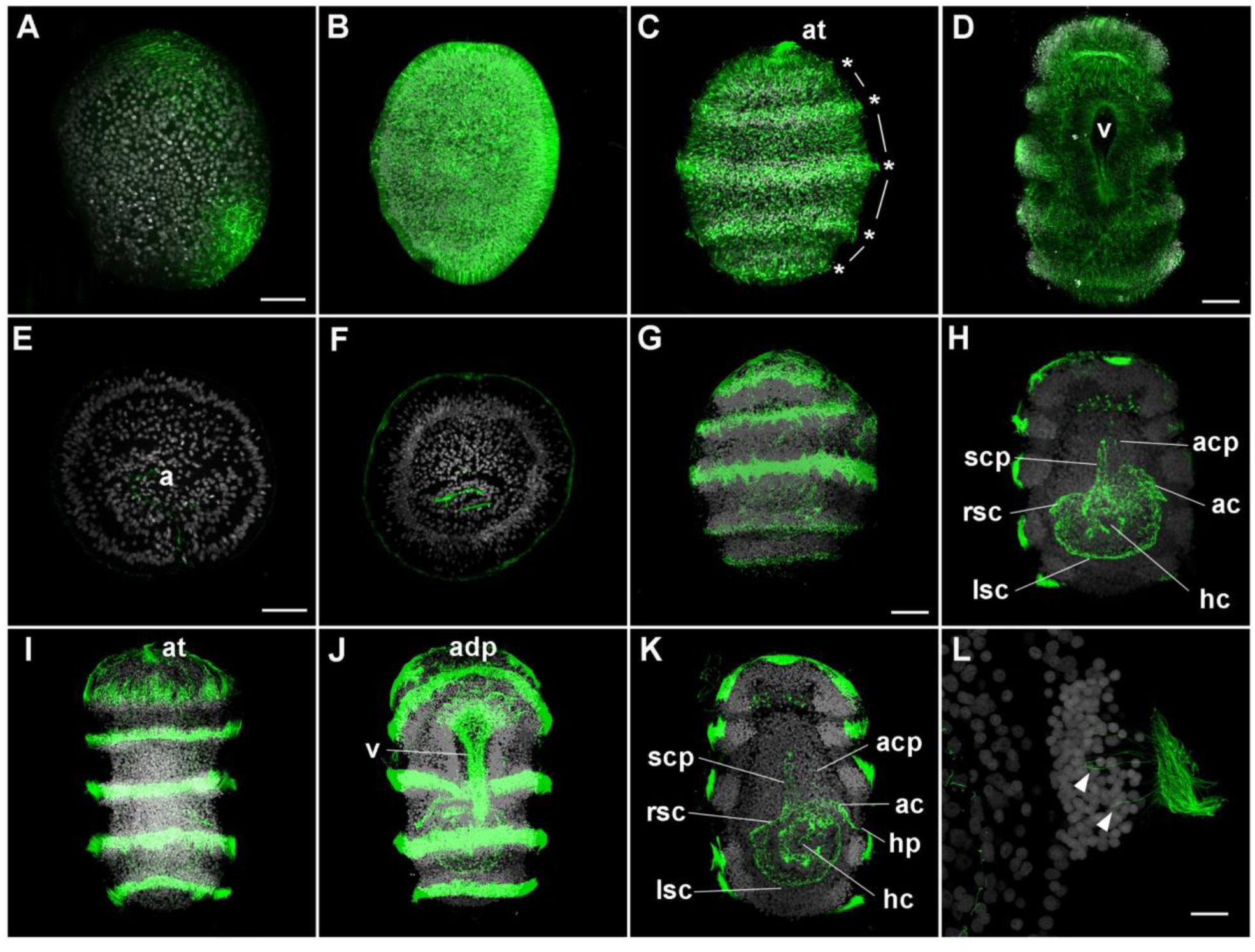
External and internal ciliogenesis in *A. mediterranea* embryos. Confocal z-projections of embryos immunolabelled with anti-ß-tubulin antibodies (**A-D**; green) or anti-acetylated α-tubulin antibodies n (**E-L**; green) and co-stained with DAPI (gray). Whole (**A**, **B**, **C**, **D**, **G**, **I** and **J**) and longitudinal sections (**E**, **F**, **H**, **K** and **L**) of representative samples. Anti-ß-tubulin marked developing cilia in 36 hpf embryo (**A**), uniformly ciliated larva (**B**), band formation (**C**) and doliolaria (**D**, ventral view) stages. Asterisks and lines in **C** indicate ciliary bands and interband cells respectively. Cilia in the ectoderm and in coelomic cavities were also immunoreactive to anti-acetylated α-tubulin: 26 hpf gastrula (**E**), uniformly ciliated larva (**F**), pre-hatching larva (**G** and **H**) and doliolaria (**I**, dorsal; **J**, ventral; **K** longitudinal view) stages; magnification of ciliated band in doliolaria larva is shown in **L**; cilia departing from inner cells are indicated (arrowhead). Legend: ac = axocoel; acp = axocoel projection; hc = hydrocoel; hp = hydropore; lsc = left somatocoel; rsc = right somatocoel; scp = somatocoel projection. Scale bars: A-K = 50 µm, L = 10 µm.

**Fig. 5.**
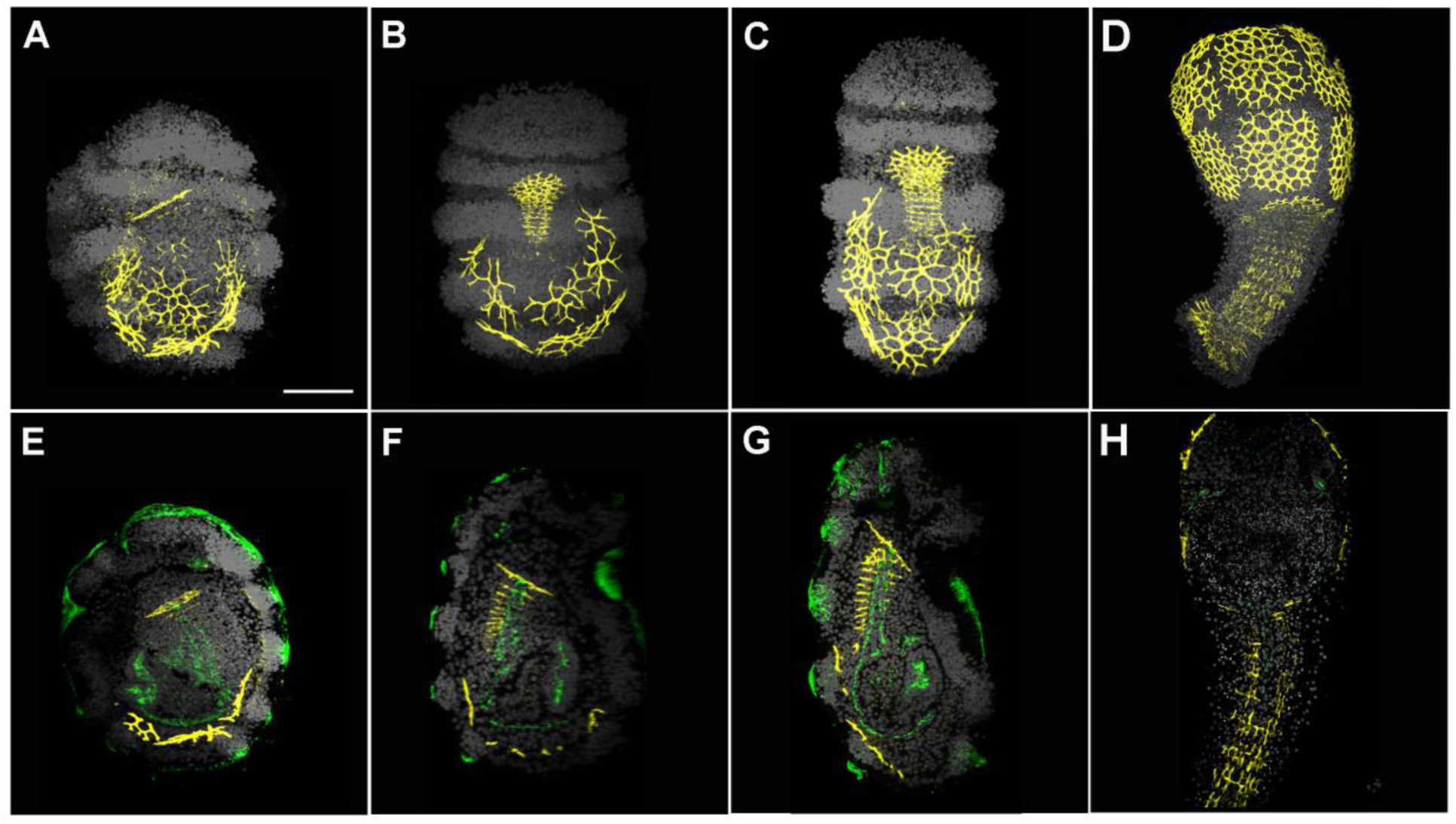
Ossicle development in *A. mediterranea*. Confocal z-projections of embryos immunolabelled with antibodies against phosphorylated SMAD1/5/8 (yellow) and acetylated α-tubulin (green) and co-stained with DAPI (grey). Whole (**A**-**D**) and mid-sagittal sections (**E**-**H**) of pre-hatching larvae, early (**A** and **E**) and late (**B** and **F**) stage, swimming doliolaria larvae (**C** and **G**) and cystidean stage (**D** and **H**). In all stages considered, the antibody labels the developing ossicles of the crinoid skeleton. Legend: ad: attachment disk; Scale bar = 100 µm.

*Band formation stage* (∼48 – 72 hpf at 17±1°C). During this developmental phase, the characteristic five ciliary bands of the doliolaria formed. At ∼48 hpf, before the development of the band-interband pattern, some morphological changes in the epithelial cells had already occurred: future band cells became thin and tightly packed to each other while interband cells remained larger (Fig. 4 C). These interbands gradually lost their cilia and the embryo epithelium turned into the typical alternation of ciliated and non-ciliated domains of doliolaria larva.

*Pre-hatching doliolaria* (∼72 –100 hpf at 17±1°C). Although still roundish in shape, the larva appeared almost completely formed. Under the fertilization membrane, the main larval features could be easily discerned, including the five transverse ciliary bands and two ventral grooves, the adhesive pit and the vestibulum (Fig. 1 G-I and Fig. 4 G). Proliferation was concentrated at the levels of the developing transverse ciliary bands and the apical tuft (Suppl Fig. 1 C), with PhH3 positive nuclei always localized on the external surface of the embryos. In internal tissues, mitotic nuclei were observed in both mesenchymal tissues and coelomic cavities (Suppl Fig. 1 D).

*Swimming doliolaria* (≥100 hpf at 17±1°C). After hatching, the doliolaria larva began swimming in the water column. The doliolaria appeared as a barrel-shaped larva 400-450 µm long. It was characterized by five ciliary bands that ran transversely along the body and a long anterior tuft emerging from the apical pit. Ventrally, two grooves could be observed, the anterior adhesive pit, used by the doliolaria to attach to the substrate at the beginning of metamorphosis, and the vestibulum covered by cilia (Fig. 1 L and Fig. 4 I and J).

### Ciliogenesis

Antibodies against acetylated α-tubulin and ß-tubulin (clone 2-28-33) were used to label cilia during embryogenesis of *A. mediterranea* (Fig. 4). Cilia first appeared during gastrulation: at 26 hpf, acetylated α-tubulin positive cilia were observed in the invaginating ento-mesodermal cells, projecting towards the developing archenteron (Fig. 4 E). Later, at ∼32 hpf, ciliogenesis began on ectodermal cells, so that at 36 hpf two undetermined areas of ciliated ectoderm, one anterior and one posterior, were present in most embryos (Fig. 4 A). At ∼48 hpf, ciliogenesis extended throughout the ectoderm, so that short and motile cilia covered all the outer surface of the uniformly ciliated embryos (Fig. 4 B). Inside, a ciliated epithelium bordered the coelomic cavities and the enteric sac (Fig. 4 F). During the band formation stage (Fig. 4 C), the change of the uniformly ciliated ectoderm into an alternation of band and interband domains occurred. This was a progressive and complex process that led to the development of the characteristic ciliated bands of the doliolaria larvae (Fig. 4 D). The band-interband pattern was already observable in pre-hatching larvae, in which cilia were strongly marked by the acetylated α-tubulin antibody (Fig. 4 G). In the swimming doliolaria, intense staining was detected in the five ciliated bands, in the apical tuft and in the ventral ciliated grove, the vestibulum (Fig. 4 I, J, Suppl. Video 1). Magnification of the band domain revealed that not all of the ciliated cells faced the external surface of the larva but some were internal with cilia projecting outside (Fig. 4 M, Suppl. Video 2). Inside the embryos, the development of the coelomic cavities proceeded and the ciliated epithelium of the recently formed hydrocoel and axocoel as well as that of the somatocoel appeared strongly labelled (Fig. 4 H, K). The axocoel branched into a thin projection directed anteriorly while the right somatocoel extended along the dorsal midline of larva.

### Endoskeleton

The elements that form the post-metamorphic skeleton are already present at the larval stage [4, 17]. We found that antibodies against phosphorylated SMAD1/5/8 (PSMAD1/5/8; Cell Signalling Technology, D5B10) specifically stains skeletal ossicles in *A. mediterranea*. While all ossicles were strongly immunoreactive across development, no cellular structure was labelled, indicating that the antibody recognizes a different epitope within the skeleton. Given this, we decided to use PSMAD1/5/8 to follow ossicle development. Pre-hatching larvae (72 hpf, Fig. 5 A and B, Suppl. Video 3) already had numerous spicules and developing ossicles: the oral and basal plates of the calyx were already recognizable while the columnar elements of the stalk were still forming. The complexity of the ossicles was highly variable among samples, but skeletal rudiments were always observed at the pre-hatching stage. In swimming doliolariae (Fig. 5 C, Suppl. Video 4), oral and basal ossicles began to acquire their typical plate shape and to form numerous stroma. Columnar stalk ossicles appeared well-developed and the rudiment of the attachment disk was observable. After metamorphosis, in the cystidean stage the skeleton was well-formed, comprising oral and basal plates, columnar stalk ossicles and the attachment disk (Fig. 5 D and H). Double staining with anti-acetylated tubulin antibody facilitated localization and characterization of the ossicles. Oral and basal plates were distributed at the periphery of doliolaria mesoderm, surrounding the coelomic and enteric cavities. The columnar ossicles were instead positioned along the dorsal midline of the larva, with the right somatocoel running at the centre of the developing skeletal structure (Fig. 5 E-G).

### Development and organization of the larval nervous system

While in *A. mediterranea* embryos the anti-acetylated α-tubulin antibody was specific for cilia, anti-ß-tubulin antibodies labelled both cilia and neural processes [4], and was employed to specifically follow neural fiber outgrowth during development (Fig. 6 A-E). Up to the uniformly ciliated larva, immunoreactivity was only observed in the cilia covering the embryo surface and bordering the internal coelomic/enteric cavities (Fig. 6 A). Starting from 48 hpf, short nerve fibers running at the base of epithelial cells were immunolabeled (Fig. 6 B), indicating the presence of maturing neurons.

**Fig. 6.**
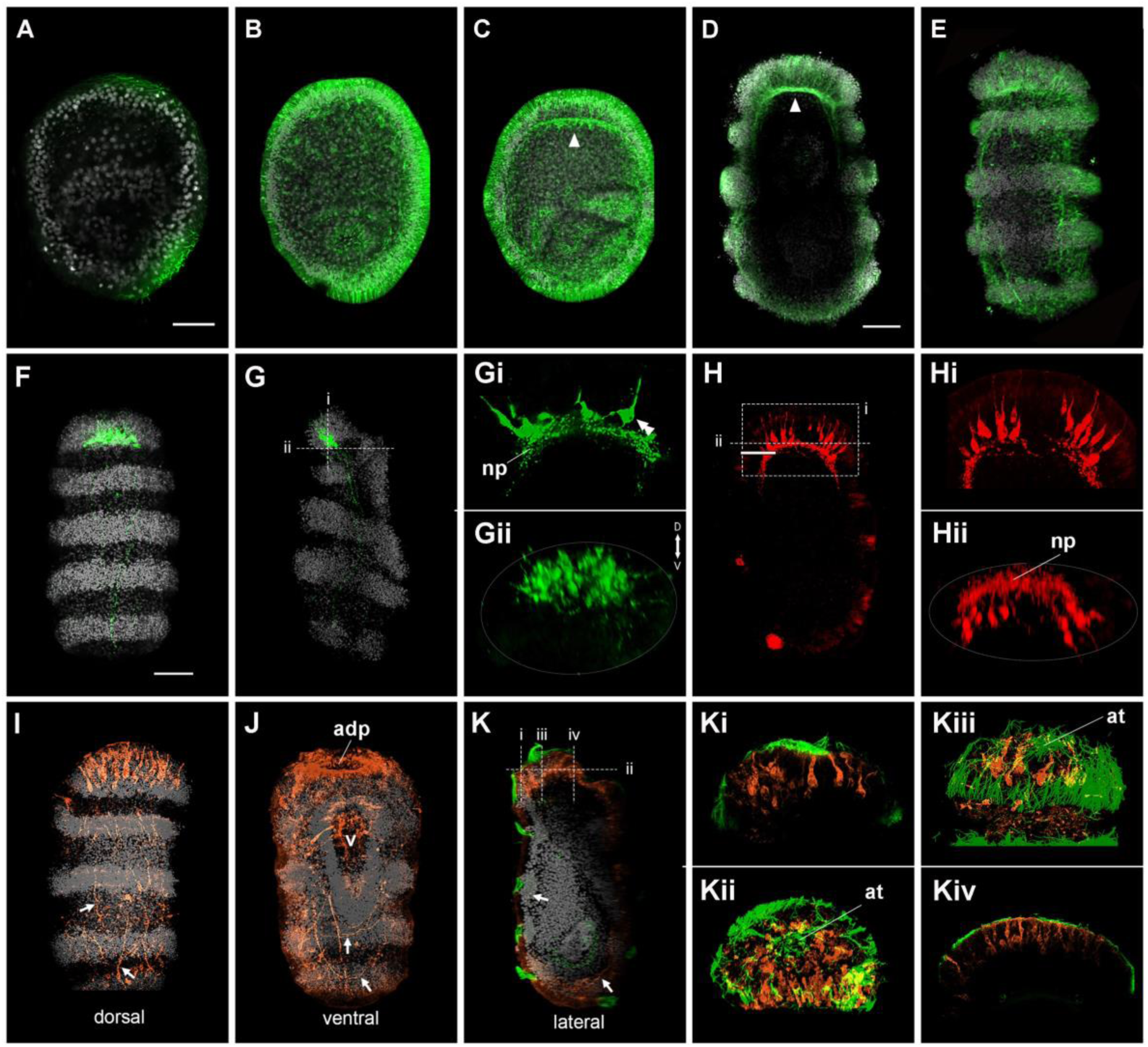
Development and cell type composition of the larval nervous system of *A. mediterranea*. Confocal z-projections of immunostained embryos and larvae showing axonal development (**A-E**) and neurotransmitter distribution (**F-K**). In all panels nuclei are stained with DAPI (grey). While neuron projections are not detected at 36hpf (**A**), anti-ß-tubulin antibodies (green) marked growing neurites in uniformly ciliated (**B**), band formation (**C**) and doliolaria (**D**, longitudinal; **E**, dorsal view) stages. The apical neural plexus is indicated with arrowhead. Serotonin (green) is localized in the anterior-dorsal apical organ and in dorsal and lateral neurites in doliolaria larvae (**F**, dorsal view; **G**, lateral view). Apical organ cells are bottle-shaped, project into the anterior plexus (**Gi**) and are concentrated dorsally (**Gii**). Double arrowheads in **Gi** indicate a single bottle-shaped cell. GABA immunoreactivity (red) was detected anteriorly (**H**, ventral view) in bottle shaped cells that projected locally in the apical plexus (**Hi**) and were located ventrally around the adhesive pit (**Hii**). Double immunostaining for glutamate (orange) and acetylated α-tubulin (green) shows glutamatergic neurons and fibers anteriorly and along the body (**I**, dorsal view; **J**, ventral view, **K**, lateral view). Arrows indicate fibers and cells of the basiepithelial neural plexus. Antero-dorsally, large cells are arranged around the apical tuft (**Ki**-**Kiii**) while ventrally small cells are organized around the adhesive pit (**Kiv**). Legend: at = apical tuft; D = dorsal; np = neural plexus; V = ventral; v = vestibule. Scale bars: 50 µm.

As development proceeded, neural processes elongated to form a basiepithelial plexus that soon became thicker in the anterior portion of the embryos (Fig. 6 C). In the swimming doliolaria, the neural plexus consisted of a complex network of nerve bundles running under the epidermis and directing towards the body surface. Anteriorly, the apical plexus consisted of densely packed fibers, from which processes connecting to the apical pit could be detected (Fig. 6 D, E).

We have previously performed immunohistochemistry on doliolaria sections showing that *A. mediterranea* has a serotoninergic apical organ [4], but its organization and the identity of neurons in the crinoid larval neural plexus remains unclear. With the newly optimized protocol for whole-mount immunohistochemistry, we examined the localization of three neurotransmitters (serotonin, GABA, glutamate) to reveal the architecture of the doliolaria nervous system (Fig. 6 F-K). Serotonin antibodies labelled a large cluster of approximatively 30 flask-shaped cells located on the anterior-dorsal side of the animal, at the level of the first ciliary band (Fig. 6 F, G). The cell bodies were located deep in the ectoderm, above the thicker portion of the neuropil, and were not directly associated with the ciliary epithelium. These serotonin-positive cells reached the anterior-dorsal surface of the embryo, where the apical pit is located, and their projections entered the anterior plexus and ran posteriorly along the dorsal and lateral sides of the larva (Figure Fig. 6 G, Gi-ii). Interestingly, serotonin-immunoreactivity starts to be detected in a smaller group of cells already at the pre-hatching stage (Suppl. Fig. 3). GABA-immunoreactive neurons were also found in the anterior portion of the larva, but were instead localized ventrally around the adhesive pit and organized in two clusters of about 8 flask-shaped cells (Fig. 6 H). The apical portion of these neurons reached the surface ectoderm between the apical and adhesive pits, while axonal projections were local and terminated in the dense anterior neuropil (Fig. 6 Hi-ii). Anti-glutamate antibodies labelled a large number of cells across the larva and were more concentrated anteriorly (Fig. 6 I-K). On the antero-dorsal surface, large flask-shaped cells surrounded the apical pit and tuft, but did not appear to have extensive axonal projections (Fig. 6 I, K, Ki-Kiii); on the ventral side, numerous small bipolar cells were distributed along the edge of the adhesive pit (Fig. 6 J, Kiv). In addition, several glutamatergic neurons were stained in the neural plexus along the larval body, both dorsally and ventrally (see arrows in Fig. 6 I, J). These cells sent projections along the anterior-posterior axis, and ventrally axons appeared to loop around the posterior end of the larva. In summary, this analysis revealed the presence of a complex nervous system in the crinoid doliolaria, characterized by multiple distinct cell types.

### Molecular patterning of the crinoid apical organ

The apical organ of eleutherozoan echinoderms is specified through a highly conserved gene regulatory network that is involved in patterning the anterior portion of the neuroectoderm [11, 38–40]. In particular, early expression of transcription factors *Six3/6* and *FoxQ2* on the animal side of the embryo defines an area called the apical plate, in which downstream genes will determine the identity of apical organ neurons. To discover whether this network is also present in crinoids, we investigated the developmental expression of *Ame_Six3/6*, *Ame_FoxQ2* and *Ame_Lhx2/9* (one of the downstream genes known to be expressed in serotonergic neurons [41]) We first identified transcript sequences for *Ame_Six3/6*, *Ame_FoxQ2* and *Ame_Lhx2/9* in an *A. mediterranea* transcriptome [42] (Suppl. Fig. 4), and then analysed their co-expression by optimizing a protocol for *in situ* HCR [43] (Fig. 7). Five developmental stages were considered: two subsequent mid gastrula stages (20 hpf and 26 hpf at 17±1°C), uniformly ciliated stage (44 hpf at 17±1°C), pre-hatching doliolaria (72 hpf at 17±1°C) and swimming doliolaria (> 100 hpf at 17±1°C; Suppl. Fig. 5). In embryos at 20 hpf, *Ame_FoxQ2* and *Ame_Six3/6* were co-expressed broadly on the anterior side of the ectoderm (Fig. 7 A). At the later mid-gastrula stage *Ame_FoxQ2* had already restricted to the animal pole, while *Ame_Six3/6* was still broad and co-expressed with *Ame_FoxQ2* in the anterior tip (Fig. 7 B) Furthermore, *Ame_Six3/6* also started to be expressed in mesendodermal cells at the tip of the archenteron cavity (Suppl. Fig. 5). In the uniformly ciliated stage, *Ame_FoxQ2* transcripts were still found at the apical tip of the embryo, but this region was now devoid of *Ame_Six3/6*, which instead formed a ring around *Ame_FoxQ2* (Fig. 7 C, D). *Ame_Six3/6* was also expressed in the anterior wall of the enterohydrocoel (Suppl. Fig. 5).

**Fig. 7.**
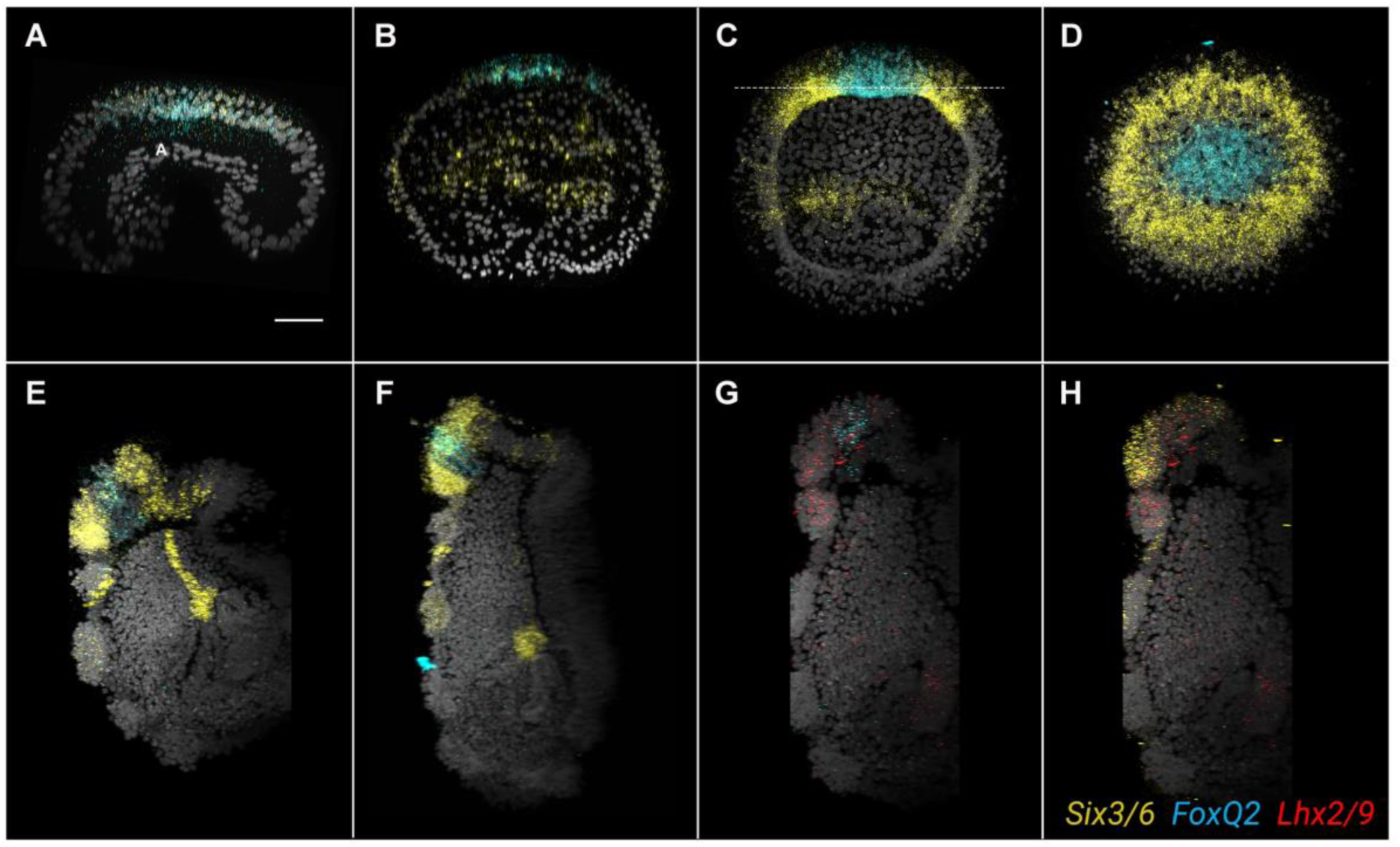
Molecular patterning of the apical region during crinoid development. Confocal images showing co-expression of *Ame_FoxQ2* (cyan), *Ame_Six3/6* (yellow) and *Ame_Lhx2/9* (red) through *A. mediterranea* development using *in situ* hybridization chain reaction. *Ame_Six3/6* and *Ame_FoxQ2* are co-expressed on the anterior/apical portion of the ectoderm at early gastrula (**A**, 20hpf) and late gastrula (**B**, 26hpf) stages, while from the uniformly ciliated stage (**C**) they form two concentric domains that persist in the pre-hatching and hatched doliolaria (**D**-**E**). At larval stages *Ame_Lhx2/9* is co-expressed with both *Ame_Six3/6* and *Ame_FoxQ2*. Scale bar: 50 µm.

In the pre-hatching doliolaria, the concentric expression of *Ame_FoxQ2* and *Ame_Six3/6* was still evident: *Ame_FoxQ2* labelled cells in the apical pit, while *Ame_Six3/6* expression was localized in the first ciliary band and in the developing adhesive pit (Fig. 7 E). Weaker *Ame_Six3/6* expression was detected in the second and third ciliary band, and a new cell population with strong *Ame_Six3/6* expression appeared just under the second ciliary band (Suppl. Fig. 5). At this stage, the downstream gene *Ame_Lhx2/9* was expressed on the dorsal side of the two ciliary bands, within the apical plate defined by *Six3/6* and *FoxQ2* (Suppl. Fig. 5). The expression of these markers remained similar in the swimming doliolaria. *Ame_FoxQ2* marked cells of the apical pit that also co-express *Ame_Lhx2/9* (Fig. 7 G). *Ame_Six3/6* transcripts were distributed in the rest of the anterior portion of the larva up to the third ciliary band, and a group of strongly-positive cells were located between the second and third band (Fig. 7 H, Suppl. Fig. 5). On the dorsal surface up to the second ciliary band, *Ame_Six3/6* was co-expressed with *Ame_Lhx2/9* (Fig. 7 H). Inside the larva, *Ame_Six3/6* labelled the axocoel, including the anterior projection of the coelom that reaches the anterior tip of the mesenchyme, while *Ame_Lhx2/9* was localized in the hydrocoel (Suppl. Fig. 5).

In other echinoderms, the network controlling anterior neuroectoderm formation is maintained on the apical side through inhibitory interactions of Wnt signalling from the vegetal side of the embryo [44] Consistently, we found *Ame_Wnt8* expressed around the blastopore and the posterior ectoderm throughout gastrulation in *A. mediterranea*, while no expression was detected at later stages (Suppl. Fig. 5).

### Development of the post-metamorphic nervous system

Recent studies have started to investigate the composition of the complex adult nervous system of eleutherozoan echinoderms [14, 45] but its developmental origin still remains elusive. Crinoids have a gradual metamorphosis in which larval tissues are rearranged to form the adult body. This feature of the crinoid life cycle could help to understand the formation of the echinoderm enigmatic adult body plan, including its nervous system. Previous studies have suggested that crinoid larval neurons degenerate at metamorphosis, and a tripartite nervous system consisting of ecto-, hypo- and entoneural components develops in the adult [16, 19]. During metamorphosis of *A. mediterranea*, we found that fibers of the neural plexus labelled by ß-tubulin were soon lost in settled larvae. The ciliary bands also disappeared after settlement, while cilia could be occasionally observed in the vestibular ectoderm (Fig. 8 A). Moreover, expression of *Ame_FoxQ2* (data now shown), *Ame_Six3/6* and *Ame_Lhx2/9* disappeared from the apical surface, which was now attached to the substrate and became the aboral side of the animal (Fig. 8 B, C). Conversely, *Ame_Six3/6* and *Ame_Lhx2/9* transcripts were still detected in the axocoel and hydrocoel respectively (Fig. 8 B, C, Suppl. Fig. 6 A-C). As metamorphosis progressed, the internal organs underwent a 90° rotation, after which the vestibule of the doliolaria reached the oral surface (corresponding to the posterior side of the doliolaria) in the cystidean stage (Suppl. Fig. 6 D-F). The ciliated hydrocoel positioned under the oral (vestibular) ectoderm, and from it five sets of three tube feet primordia emerged and were visible using antibodies to acetylated α-tubulin (Suppl. Fig. 6 D). Moreover, the post-metamorphic nervous system started to develop: antibodies to ß-tubulin labelled a prominent stalk nerve already in cystidean samples, indicating that the entoneural system differentiated shortly after metamorphosis (Fig. 8 D). In late cystidean stages, fibers also directed orally from the aboral nerve centre (Fig. 8 D). Interestingly, already in the cystidean stage the oral ectoderm started to express both *Ame_Six3/6* and *Ame_Lhx2/9* at the level of the tube feet primordia (Fig. 8 E, F).

**Fig. 8.**
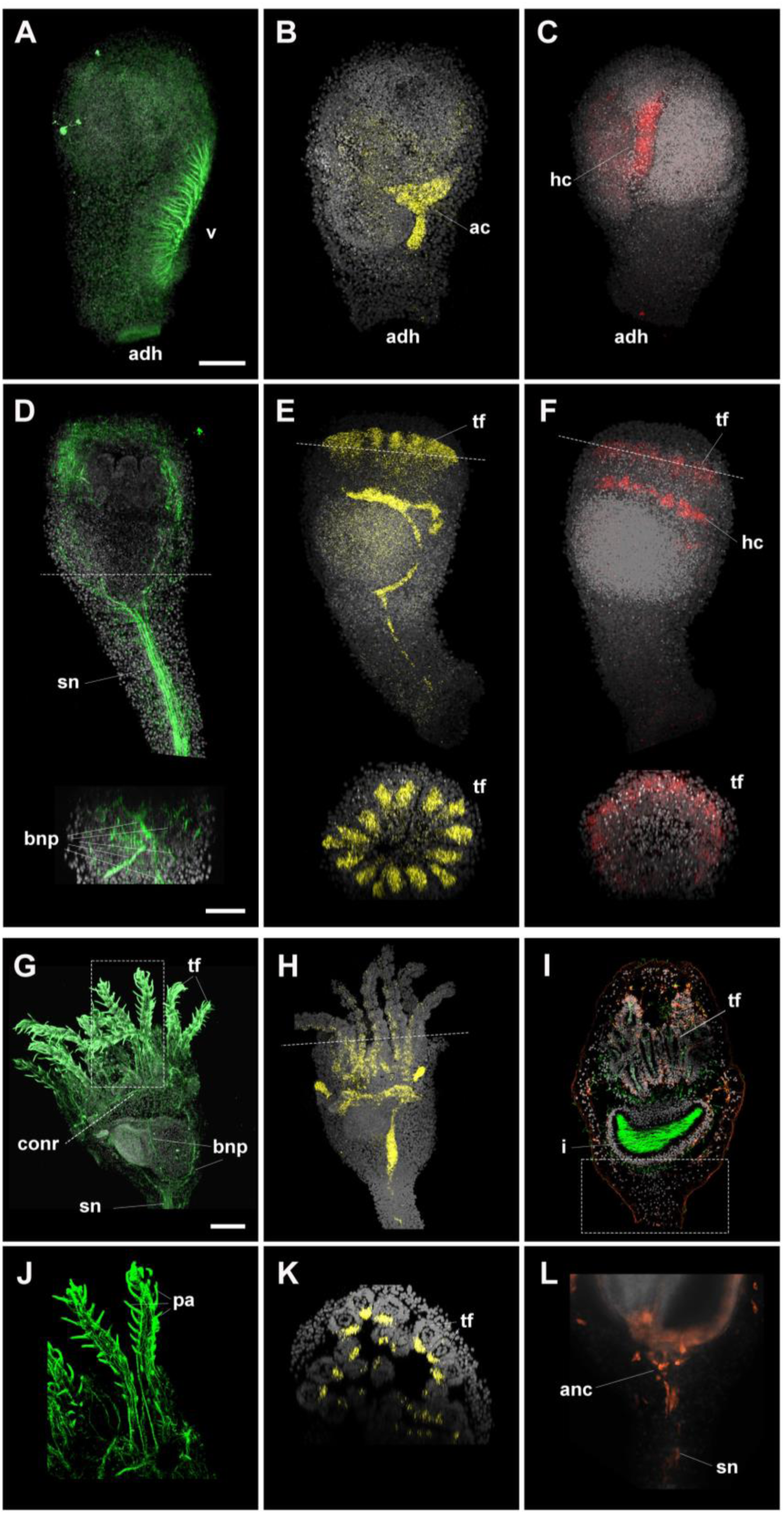
Formation of the post-metamorphic nervous system in *A. mediterranea*. Confocal z-projections immunolabelled with anti-ß-tubulin (green) (**A**, **D**, **G**, **J**), anti-glutamate (orange) and anti-acetylated α-tubulin (green) antibodies (**I**, **L**) or labelled with probes against *Ame_Six3/6* (yellow) (**B**, **E**, **H**, **K**) and *Ame_Lhx2/9* (red) (**C**, **F**). Signal for all markers disappears in the apical region after settlement (**A**-**C**). In the cystidean stage (**D**-**F**) the post-metamorphic nervous system forms: neurites are labelled with ß-tubulin and the tube feet ectoderm starts to express *Ame_Six3/6* and *Ame_Lhx2/9*. Transverse sections for each image are shown at the level indicated by the dashed lines. In the pentacrinoid stage (**G**-**L**) ß-tubulin marks neural fibers in the stalk nerve, radial nerve primordia, circumoral nerve ring and tube feet, including the papillae (**G**, **J**); *Ame_Six3/6* is expressed in the oral side of the tube feet ectoderm (**H**, **K**); and glutamate is localized in the tube feet, neurons of the epidermal plexus, aboral nerve centre and stalk nerve, while acetylated α-tubulin shows cilia in the hydrocoel of the tube feet, somatocoel and intestine (**I**, **L**). J, K and L panels show specific portions of the pentacrinoid indicated by dashed lines/boxes. Legend: ac = axocoel; adh = adhesive pit; anc = aboral nerve centre; bnp = brachial nerve primordia; conr = circumoral nerve ring; hc = hydrocoel; i = intestine; pa = papillae; sn = stalk nerve; tf = tube feet; Scale bars: 50µm

It has been shown previously that the pentacrinoid nervous system comprises a complex basiepithelial net of serotonergic, GABAergic and peptidergic neurons, concentrated at the level of the tube feet, and a conspicuous entoneural system comprising cholinergic and peptidergic cells [4, 27]. Immunoreactivity for ß-tubulin showed that the post-metamorphic system was highly interconnected, with five bundles of fibers directing from the entoneural aboral nerve centre to a circumoral nerve ring, which was in turn connected to fibers that run in the tube feet (Fig. 8 G, J). Curiously, at this stage the tube feet expression of *Ame_Six3/6* persisted in the later pentacrinoid stage, but was restricted to the internal portion of each tube feet, facing the mouth (Fig. 8 H, K). Here we further expanded this analysis by characterizing the localization of glutamatergic neurons (Fig. 8 I, L). A large number of positive cells were detected in the tube feet (Fig. 8 I, Suppl. Fig. 6 G). Contrary to serotonergic neurons, these cells did not form an obvious axonal net, but positivity was found mainly in the cell bodies. Interestingly, positive puncta were also found at the tip of the papillae (Suppl. Fig. 6 G). Scattered glutamatergic neurons were also labelled across the calyx epidermis (Fig. 8 I). These cells appeared to be more concentrated on the aboral side and possess short axonal projections. A variable number of neurons were also labelled in the entoneural system (Fig. 8 L). A few positive cells were sometimes visible just oral to the aboral nerve centre, while several immunopositive fibers and scattered cell bodies could be seen along the stalk nerve (Fig. 8 L, Suppl. Fig. 6 G).

## Discussion

Although crinoids occupy a key phylogenetic position as the sister group to the other echinoderms [6], they have been generally neglected in modern developmental biology, and the molecular mechanisms regulating their embryogenesis have remained largely unexplored. The difficulty in collection and maintenance under laboratory conditions of many crinoid species, the scarce information about cues governing spawning and reproduction, and the low availability of techniques to investigate the molecular control of development have together hampered crinoid employment in evo-devo studies [16, 17, 28–30]. In this work, we show how to overcome most of these obstacles, as *A. mediterranea* can be found in shallow waters during its reproductive season, adults can be easily collected, and gamete release could be induced in aquaria to obtain hundreds of fertilized eggs. Although *A. mediterranea* is an external brooding species [33], and embryos normally develop attached to female’s genital pinnules [4, 17], our method allowed us to detach embryos from female individuals and successfully rear them *in vitro* to precisely follow embryogenesis. Furthermore, we have optimized techniques for whole-mount immunohistochemistry and multiplex *in situ* HCR that allowed us to monitor development and characterize embryonic, larval and post-metamorphic structure to better understand the body plan of this enigmatic group of echinoderms.

### Crinoid embryogenesis and formation of larval structures

Our current understanding of crinoid early development is based on old microscopy studies from the late nineteenth century and early twentieth century [15, 16]. These works reported detailed descriptions of the cellular and subcellular modifications happening during embryo development [34–36], but standardized developmental timing and staging under controlled laboratory conditions was lacking. Thus, we followed *A. mediterranea* embryogenesis *in vitro* and described each embryonic phase together with its developmental timing using high-resolution images. Interestingly, we found no differences in survival, embryo morphology and developmental timing between embryos developing *in vitro* and on females’ pinnules, suggesting that maternal contribution after spawning might be limited to preventing predation and maintaining seawater clean and well oxygenated. Consistent with previous descriptions of crinoid development, we found that in *A. mediterranea* cleavage was holoblastic, radial and unequal, forming smaller animal blastomeres and larger vegetal blastomeres from the 8-cell stage. Cell divisions were asynchronous and occurred faster in the animal blastomeres, often making embryo morphology irregular and divisions difficult to follow. Curiously, we also observed that embryos of the same batch developed asynchronously both *in vitro* and on genital pinnules. These features were not related to our experimental conditions, as they have been already reported [15, 46], and they appear to be a specific characteristic of feather star development. Earlier microscopy works reported that in crinoid species gastrulation began as unipolar ingression of single cells before gradually turning into an invagination process [15, 16]. This was refuted in *Florometra serratissima* and in *A. japonica*, for which gastrulation by invagination [47, 48] or by holoblastic involution [46] were proposed instead. According to our data, *A. mediterranea* development proceeds with gastrulation by invagination, preceded by shape changes of vegetal pole cells. These blastomeres gradually acquire a flask-like morphology, likely narrowing at the apical side. Apical constriction is a well-known mechanism that drives primary invagination in both sea urchin and amphibians [49–51], and might therefore be functional to the bending of the ento-mesodermal layer in *A. mediterranea*. During gastrulation, the thick ento-mesodermal layer quickly reaches the animal pole, in parallel with a drastic reduction of the blastocoel size, which is progressively filled by mesenchymal cells as previously shown in other crinoids [46, 47]. This is in contrast with the initial phases of gastrulation in sea urchin, in which the ento-mesoderm only occupies a small portion of the embryo cavity [49]. The increase in embryo complexity during and after gastrulation is accompanied by widespread proliferation. A peculiar pattern of cell division was observed in the ectoderm during gastrulation, where nuclei of dividing cells were concentrated on the apical surface. This condition is similar to the interkinetic nuclear migration observed in the vertebrate neural plate and in other pseudostratified epithelia [52] in which nuclei migrate along the apical-basal length across the cell cycle.

One of the hallmarks of echinoderm larvae is the presence of ectodermal ciliary bands. Previous studies have highlighted that in many crinoid species the ciliary bands of the doliolaria form from an initial uniformly ciliated ectoderm [17, 24, 48]. In *A. mediterranea,* both ento-mesodermal and ectodermal tissues are highly ciliated. The formation of ectodermal cilia started from about 32 hpf in two domains, one located near the animal pole and one more posterior, and then expanded through the ectoderm, so that by 48 hpf uniformly ciliated larvae start to rotate inside the fertilization membrane. This mechanism of cilia formation occurred at the same stage and with similar dynamics to another feather star, *F. serratissima*, where the first ectodermal motile cilia form at the animal pole and then progressively expand through the embryo surface [48]. As described in *Florometra*, in *A. mediterranea* the band-interband pattern of the doliolaria epidermis formed as a result of a combination of proliferation events and morphological changes in cell shape and arrangement. In the developing ciliary bands, cells became smaller and tightly packed, probably due to the extensive proliferation observed in this area; only a few dividing cells were instead detected in the interband domains where cells remained large and lost their cilia. The pre-hatching larva already showed well-developed transverse ciliary bands, apical tuft and two ventral ciliated grooves, the vestibulum and the adhesive pit.

We found the use of antibodies against acetylated α-tubulin and PSMAD1/5/8 to be instrumental in describing the formation of internal structures during larval development, including the coelomic and enteric cavities and the skeleton. After invagination, a ciliated epithelium bordered the archenteral sac, which divides into an anterior entero-hydrocoel and a posterior somatocoel in the uniformly ciliated stage. Then, the entero-hydrocoel elongated acquiring a dumbbell-like shape while the somatocoel curved around it, assuming a crescent shape. Finally, by the pre-hatching doliolaria all coelomic cavities and the enteric sac are formed. From this stage, acetylated α-tubulin staining also highlighted the presence of two anterior coelomic projections: one was part of the right somatocoel and was located within the rudiments of the stalk columnar ossicles, while the other branches from the axocoel, was difficult to visualize by microscopy staining [4] and almost reached the anterior ectoderm. Interestingly, already at the pre-hatching stage several spicules and developing ossicles were recognizable. Rudiments of the future oral and basal plates of the post-metamorphic calyx were observed in the posterior portion of the larvae, surrounding the coelomic and enteric cavities, while columnar ossicles developed along the dorsal midline within the mesenchyme, surrounding the somatocoel projection. Primordia of the skeletal plates have been previously found in doliolaria larvae of different feather star species [48, 53, 54]. A high variability in ossicle number and complexity was previously reported in *F. serratissima* larvae, in which ossicles first appeared variably between initial and late doliolaria [53]. Conversely, in *A. mediterranea* numerous ossicles were already present in pre-hatching larvae, suggesting that skeletogenesis began early during development. Taken together, these results complement previous descriptions of coelomic and skeletal development in other crinoid species [15, 16] and highlight the differences between feather stars and sea lilies [15, 16, 21, 21, 34–36, 53], indicating that immunohistochemistry is a valuable tool to understand the development of crinoid tissues.

### Cell type diversity and organization of the crinoid larval nervous system

Previous studies highlighted the presence of serotonergic neurons in the anterior ectoderm of crinoid larvae [4, 17, 19]. However, technical limitations of existing protocols, mostly restricted to tissue sections and without the possibility to test gene co-expression, together with the scant data on embryogenesis, have impeded the description of the early development and of the three-dimensional organization of the larval nervous system. Here, using newly optimized protocols for whole mount immunohistochemistry and *in situ* HCR, we provided a more comprehensive characterization of the crinoid larval nervous system and its molecular specification. Short neural fibers were first detected at the base of the epidermis at 48 hpf and later became an intricate basiepithelial plexus. In the pre-hatching doliolaria, the nervous system consisted of a network of nerve bundles running under the epidermis and thicker in the anterior region, where serotonergic neurons were already visible, consistent with previous data showing localization of serotonin in uniformly ciliated embryos of sea lily [19].

In the swimming doliolaria, the larval nervous system was fully differentiated and consisted of a complex network of serotonergic, GABAergic and glutamatergic cells (Fig. 9 A). A prominent apical organ was located in the antero-dorsal region and composed of ∼30 serotonergic neurons that sent long-ranging projections posteriorly through the anterior neuropil. While the connectivity of serotonergic cells in other echinoderms is still unclear, our results suggest that these neurons connect with other parts of the neural plexus to direct larval activities. The number of serotonergic cells varies greatly across echinoderms: in sea urchins, early pluteus larvae have 4-6 serotonergic cells, although their number increases in later stages [55]. Conversely, the starfish bipinnaria and the holothuroid auricularia larvae already have 30-50 and ∼20 serotonergic neurons respectively [11, 56–59]. This variability indicates that while serotonergic neurons are a highly-conserved feature of larval development, their number and distribution evolved independently in each echinoderm lineage. In addition to serotonergic cells, around the *A. mediterranea* apical pit multiple large glutamatergic cells were present, which could represent support cells often associated with apical organs [60]. Conversely, the adhesive pit was bordered by large GABAergic cells that sent local projections to the anterior neuropil, and numerous small glutamatergic cells. If serotonergic neurons of the apical organ have a sensory function, as described in other ciliated larvae [60], then it is possible that they interact with GABAergic neurons to direct settlement and metamorphosis. In fact, GABA has been previously found to be involved in metamorphosis in sea urchin larvae [61]

**Fig. 9.**
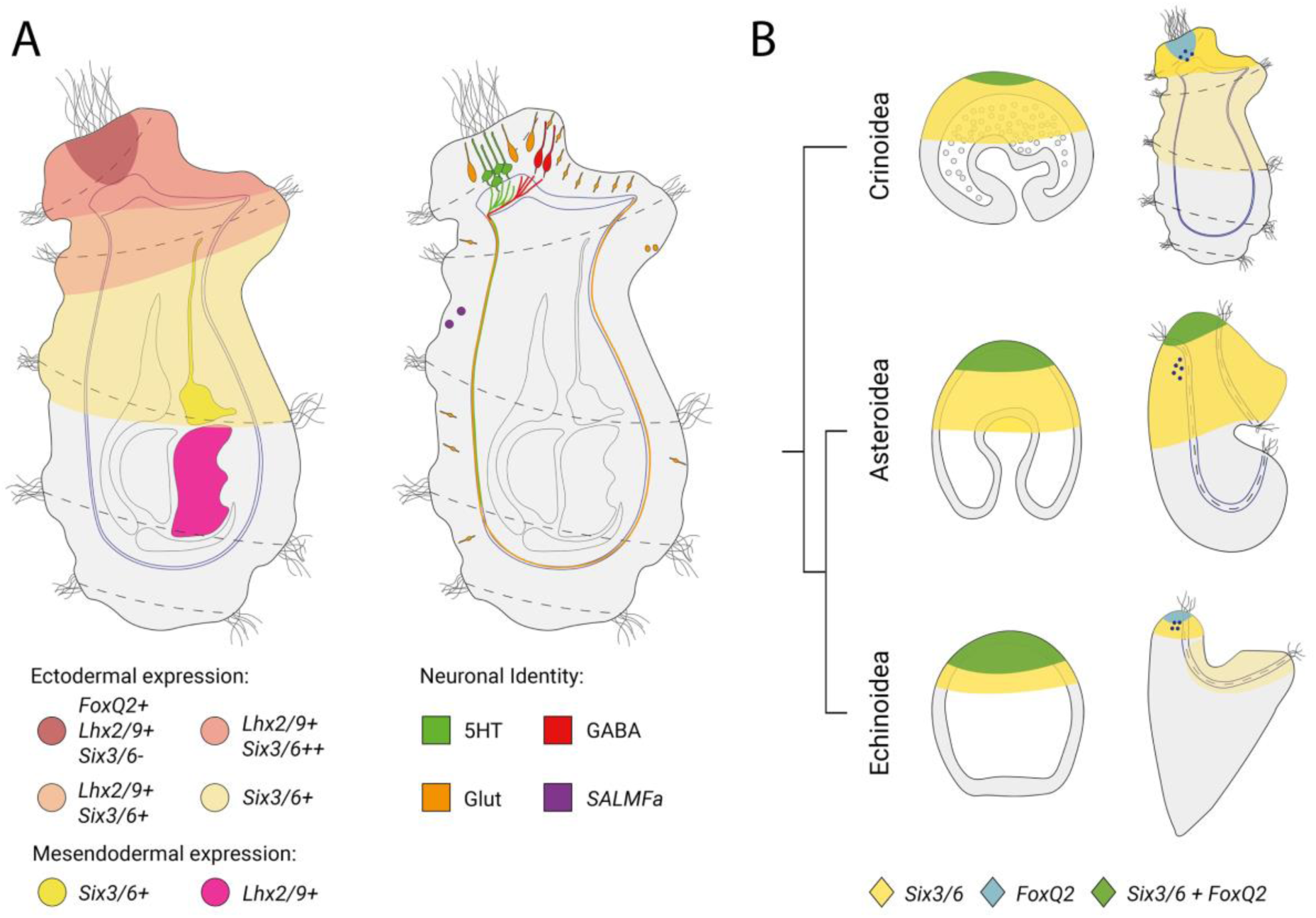
Summary diagram of the crinoid larval nervous system. **A**) Expression data and neural populations in the doliolaria larva (lateral view). **B**) comparison of expression of *FoxQ2* and *Six3/6* across echinoderms.

While the apical organ has been previously identified in crinoids, neurons in the basiepithelial plexus remained uncharacterized. In addition to serotonergic projections from the apical organ, we found that the diffuse nerve plexus contained several glutamatergic neurons and fibers, organized both ventrally and dorsally. To our knowledge, this represents the first insight into the identity of crinoid basiepithelial neurons, and suggests that the doliolaria nervous system has a large glutamatergic (likely excitatory) component. Contrary to most other echinoderm larvae studied to date, the crinoid plexus does not appear to be associated with the ciliary bands; instead, the nerve plexus is broadly distributed and axons travel through the entire ectoderm.

### Conservation of apical organ development

The molecular control of apical organ development has been thoroughly investigated in multiple echinoderms including sea urchin and starfish [41, 62, 63]. Many transcription factors acting through gastrulation and organogenesis have been shown to be required for the correct specification and positioning of the apical organ. Of these, *Six3/6* and *FoxQ2* are considered to be upstream in the regulatory interactions, as experimental knock-down of both genes lead to the loss of anterior serotonergic neurons, as well as of downstream transcription factors, such as *Fezf*, *Nk2.1* and *Lhx2/9* [41, 64–66]. Comparative studies on the expression of these genes in multiple phyla have recently led to the hypothesis that a conserved anterior gene regulatory network (aGRN) might be involved in the specification of the anterior neuroectoderm across Bilateria [38, 39, 62, 67]. However, there are also significant differences in the organization of serotonergic neurons and the expression of aGRN transcription factors between echinoderm classes. For example, in starfish serotonergic neurons are born throughout the apical surface of the larva and then migrate to form a dorsal bilateral structure within the ciliary band, while in sea urchins serotonergic cells develop locally dorsal to the ciliary band [11, 55, 56]. Moreover, in starfish *FoxQ2* and *Six3/6* are co-expressed in the apical plate throughout development, while in sea urchin after early co-expression the two genes form mutually-exclusive concentric rings of expression [41, 64] (Fig. 9 B). Previous chromogenic *in situ* analysis in sea lily and feather star larvae showed that *Six3/6* is expressed in the anterior portion of the ectoderm [20, 22]. To explore the conservation of apical organ development in crinoids, we produced the first instance of multiplex fluorescent *in situ* hybridization in crinoids, showing that co-expression of *Ame_FoxQ2* and *Ame_Six3/6* is already visible during gastrulation in a broad anterior ectodermal domain (apical plate) and then restricts anteriorly. After gastrulation, *Ame_Six3/6* disappeared from the *Ame_FoxQ2* domain and formed a ring around the anterior tip of the animal, as in sea urchin. After the formation of the apical plate, the downstream gene *Ame_Lhx2/9*, which is required for the formation of serotonergic neurons in starfish [41], started to be expressed within the first two ciliary band and was co-expressed with *Ame_FoxQ2* anteriorly, in the area where serotonergic neurons form. Taken together, these results support the conservation of the aGRN in crinoids, and indicate that the ancestral organization of the network in echinoderms included the formation of concentric rings of *Six3/6* and *FoxQ2* expression in the apical plate.

### The crinoid post-metamorphic nervous system

The adult nervous system of echinoderms follows the pentaradial symmetry of the body, but its origin and early development have long remained obscure [68]. In crinoids most of the larval tissues are maintained in the post-metamorphic stages, but the doliolaria nervous system degenerates at metamorphosis, as we demonstrated with the loss of apical neurites and of the apical expression of aGRN genes shortly after settlement. We have previously shown that in the post-metamorphic pentacrinoid stage the nervous system already comprises ectoneural and entoneural components [4]. Here, we further revealed that these neurons begin to form already at the cystidean stage, in which neurites are visible at the level of the developing tube feet and in the entoneural stalk nerve, aboral nerve centre and brachial nerve primordia. Moreover, we also discovered that the early tube feet ectoderm, where a concentrated net of serotonergic, GABAergic and glutamatergic neurons develops in the pentacrinoid [4], starts to express the anterior markers *Ame_Six3/6* and *Ame_Lhx2/9* already from the cystidean stage. A recent study has reported that the tube feet ectoderm of the *A. japonica* cystidean additionally expresses *Otx* and *Pax4/*6 [22]. Taken together, these results indicate that the newly-formed post-metamorphic ectoneural system in crinoids expresses conserved anterior markers, consistent with what was recently discovered in eleutherozoans [45]. The inactivation of these genes during settlement and re-activation in the cystidean stage, however, suggests that this network was secondarily deployed in the adult stage during echinoderm evolution.

## Conclusions

The echinoderm body plan has long puzzled zoologists [16]. In order to understand the evolution of this phylum, and more broadly of deuterostomes, it is essential to compare data from a variety of taxa. Although crinoids are in an ideal phylogenetic position to investigate the evolution of this enigmatic phylum, biological and technical problems have hindered their study, resulting in far less information available for this group compared to other echinoderm classes. In this work, we developed a successful method for culturing *A. mediterranea* embryos *in vitro* and provided a standardized timetable of embryogenesis, making early crinoid embryos accessible to many experimental approaches. Our analysis also represents the most comprehensive description of the crinoid larval nervous system to date, revealing a complex organization and the presence of a large number of morphologically and functionally diverse cell types. These results support the conservation of an aGRN controlling the formation of the apical organ in ambulacrarians (echinoderms and hemichordates). Overall, our work will contribute to promote the employment of crinoids in developmental biology and comparative studies, allowing the comparison of crinoid development with other well-established models to help clarify many unsolved questions on deuterostome evolution.

## Methods

### Animal collection and maintenance

Adults of *A. mediterranea* were collected during winter 2021 at La Spezia Gulf (Ligurian Sea, Italy). Animals were immediately transferred to the laboratory, where they were maintained in aquaria filled with seawater (SW) and provided with closed multi-circulation system as well as mechanical, chemical and biological filters. Aquaria physical and chemical parameters were regularly checked and adjusted if necessary [69]. Temperature was set at 17±1°C, and natural light conditions were preferred. Animals were fed with commercial feed for seawater filter-feeding animals (Coralific Delite™) and their general health condition was monitored daily.

### Embryo cultures

In captivity, mature animals of both sexes were kept in the same tank and were found to respond to stressors by spawning; water turbulence and/or intense light (torch) appeared to be the main cues. Generally, male individuals spawned first, possibly stimulating mature females to release the eggs that were fertilized and retained on genital pinnules. To detach fertilized eggs from genital pinnules, gravid females were stressed, 15 min after spawning, by transferring them into 100 ml glass crystallizers filled with SW, and were left resting for half an hour with occasional pipetting of water on their pinnule surfaces. Released zygotes were then collected and transferred from crystallizer bottoms to sterile plastic petri dishes (100 embryos/dish), any impurities were removed and embryos were reared in filtered SW at 17±1°C until doliolariae hatched. This procedure allowed the collection of hundreds of embryos, but many other remained attached on female genital pinnules. Females were moved back to the aquaria and embryonic development on pinnules was daily monitored under a stereomicroscope until hatching. Embryos maintained *in vitro* were regularly checked as well, dead specimens were promptly removed, and optimal culture conditions were maintained by total water replacement on a daily basis. *In vitro* hatching rate was calculated for three batches of 100 embryos each as: (number of hatched larvae / initial zygotes) x 100. During the whole culture period, embryos at different developmental stage were fixed for the following analyses (see each section for details on fixation). During the first 12 hpf, embryos were fixed every 30 min; subsequently, specimens were collected every hour between 20 and 36 hpf and at 44, 48, 60, 72, 96 and 100 hpf.

### Whole mount phalloidin staining

To study cleavage, embryos from 1 hpf to 12 hpf were processed for phalloidin staining. After fixation in 4% paraformaldehyde, 0.5 M NaCl, and 0.1 M 3-(N-morpholino)propanesulfonic acid (pH 7.5; MOPS fixative) for 90 min at room temperature, samples were washed in phosphate buffered saline + 0.01% Triton X (PBST) and incubated in 1.5% H_2_O_2_, 5% formamide and 0.2xSSC solution for 15 min. Permeabilization was performed in 1% DMSO in PBST for 15 minutes. Embryos were then washed in PBST and incubated for 2 h in 50% PBST/50% inactivated goat serum (NGS). Finally, samples were stained at 4°C for two days with Phalloidin Atto 488 (Sigma-Aldrich), diluted 1:50 in PBST, washed several times in PBST, mounted in 80% glycerol on microscope slides and examined with a Leica SP8 or a Nikon A1 laser scanning confocal microscope.

### Whole-mount immunolocalization

In previous work, we designed a protocol for immunofluorescence on paraffin sections of doliolaria larvae [4]. Here, we optimized a new protocol for whole-mount immunofluorescence in different developmental stages, across *A. mediterranea* embryogenesis. Embryos from the mid gastrula stage (20 hpf) to hatching doliolaria were fixed in MOPS fixative and stored in 100% methanol at −20°C. Samples were washed with PBST and bleached in 1.5% H_2_O_2_, 5% formamide and 0.2xSSC. Bleaching was performed at room temperature under light for 35 min for embryos (up to the pre-hatching stage) and 45 min for larvae (pre-hatching stage and doliolaria). Embryos were permeabilized by washing them with a permeabilization solution (1% DMSO in PBS + 1% TritonX) for 10 min and then left overnight in PBST at 4°C. Larvae were instead left overnight in permeabilization solution at 4°C, then further permeabilized with 2 µg/ml proteinase K for 10 min at 37°C and post-fixed in MOPS fixative for 30 min. All samples were preincubated in 50% NGS in PBST for 2 hours and incubated in the same solution with primary antibody (Table 1). After several washings in PBST, they were preincubated in 1% BSA in PBST and incubated overnight at 4°C in PBST with the corresponding secondary antibody (Table 1) and 4′,6-diamidino-2-phenylindole (DAPI) (1:1000) to mark the nuclei. After PBST washings, samples were mounted with DABCO or 80% glycerol on microscope slides and observed using a Nikon A1 laser scanning confocal microscope or an Olympus V3000 laser scanning confocal microscope.

**Table 1.**
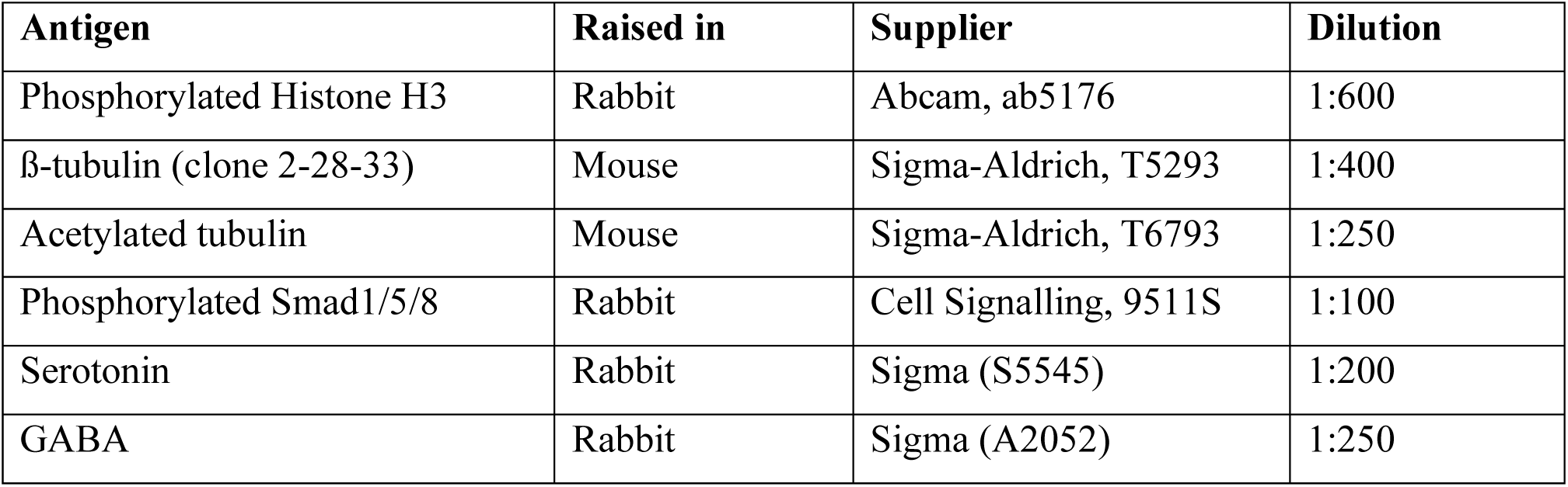
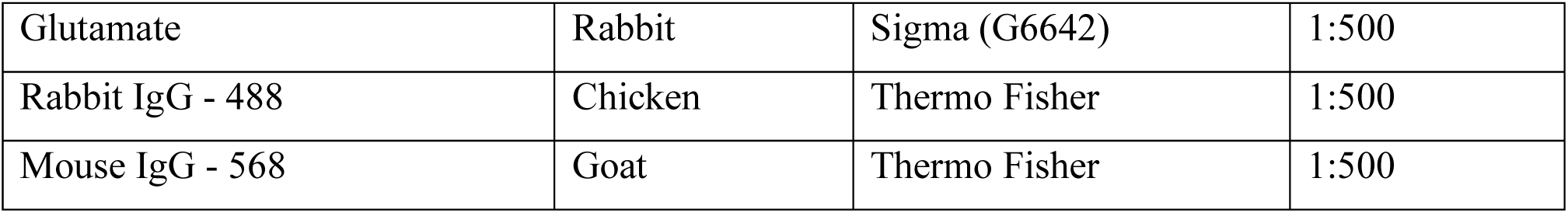
List of antibodies and their dilution used in this study.

| Antigen | Raised in | Supplier | Dilution |
| --- | --- | --- | --- |
| Phosphorylated Histone H3 | Rabbit | Abcam, ab5176 | 1:600 |
| β-tubulin (clone 2-28-33) | Mouse | Sigma-Aldrich, T5293 | 1:400 |
| Acetylated tubulin | Mouse | Sigma-Aldrich, T6793 | 1:250 |
| Phosphorylated Smad1/5/8 | Rabbit | Cell Signalling, 9511S | 1:100 |
| Serotonin | Rabbit | Sigma (S5545) | 1:200 |
| GABA | Rabbit | Sigma (A2052) | 1:250 |
| Glutamate | Rabbit | Sigma (G6642) | 1:500 |
| Rabbit IgG - 488 | Chicken | Thermo Fisher | 1:500 |
| Mouse IgG - 568 | Goat | Thermo Fisher | 1:500 |

### Whole mount *in situ* hybridization

Transcript sequences of *Ame_Six3/6*, *Ame_FoxQ2* and *Ame_Lhx2/9* retrieved from the *A. mediterranea* transcriptome [42] using the reciprocal best hits BLAST approach (Suppl. Table 1). Then gene orthology was tested with phylogenetic analysis using SeaView [70]. Protein sequences were aligned using MUSCLE [71] and phylogenetic tree was constructed using the Neighbour Joining method with 1000 bootstrap repetitions.

Chromogenic *in situ* hybridization was performed as previously described [4]. Digoxigenin labelled riboprobes specific for *Ame_FoxQ2* (Fwd: GCAATGGCAATTATGAACTCACCA; Rv: CCGTTAGCGCTTCGGCTTAT) were synthetized as in [72].

For *in situ* HCR, probes for *Ame_Six3/6*, *Ame_FoxQ2* and *Ame_Lhx2/9* were ordered from Molecular Instruments®. A whole mount HCR protocol was optimized in embryonic, larval and post-metamorphic stages of *A. mediterranea* by combining aspects of an amphioxus HCR protocol [73] with the chromogenic *in situ* hybridization protocol previously designed for this crinoid species [4]. All samples were rehydrated through a methanol/H_2_O series in Eppendorf tubes and washed in PBST. Pentacrinoids were decalcified overnight in a solution of 5% EDTA in nuclease free water (nfH_2_O). All samples were bleached in bleaching solution on aluminum foil. Doliolaria larvae were bleached for 45 minutes, while embryonic and pentacrinoid stages were bleached for 20 minutes. Doliolaria larvae were incubated in permeabilization solution overnight at 4°C; while embryonic and pentacrinoid stages were incubated for 15 and 30 minutes, respectively, and then left in PBST overnight at 4°C. The following day, doliolaria larvae were further permeabilized with PK treatment at a concentration of 4 μg/ml at 37°C for 8 minutes followed by post-fixation in 4% PFA. All stages were pre-hybridized in hybridization buffer for 2 hours at 37°C and then incubated with probes in hybridization buffer for 5 days at 37°C. After 6 days washed in wash buffer and several in SSCT, samples were left in amplification buffer for 30 minutes and then incubated for at least 20 hours at room temperature in the dark with the hairpins. All samples were washed in SSCT the following day and incubated overnight with PBST + 1μg/ml DAPI at 4°C in the dark. Finally, samples were mounted in 100% glycerol in glass-bottomed dishes and imaged using an Olympus V3000 inverted confocal scanning microscope.

### Light Microscopy

Embryos from 8 hpf to 48 hpf were processed for standard light microscopy as previously described [74]. Briefly, samples were dehydrated through an ethanol series and embedded in Technovit resin (Heraeus Kulzer) according to manufacturer’s guidelines. Sections of 5 μm were cut with a microtome and stained with hematoxylin and eosin. Samples were observed under a Leica microscope and photographed using a Leica DFC-320-C camera.

### Scanning electron microscopy (SEM)

SEM analyses were performed as described in [75]. Briefly, zygotes and doliolaria larvae were fixed in 2% glutaraldehyde in SW for 2 hours and washed overnight in filtered SW at 4°C. Samples were post-fixed with 1% OsO_4_ in SW and glucose for 2 hours, washed in distilled water, and dehydrated. Absolute ethanol was gradually substituted with hexamethyldisilazane (Sigma-Aldrich). Samples were left to dry, mounted on stabs with carbon adhesive discs, then Au sputtered using a Bal-Tec SCD 050 Sputter Coater (Bal-Tec AG, Balzers, Liechtenstein), and examined with a scanning electron microscope (LEO1430).

## Supporting information

MercurioGattoni_SupplementaryFigures

## List of abbreviations

aGRN: anterior gene regulatory network
hpf: hours post fertilization
SEM: scanning electron microscopy
PhH3: phosphorylated histone 3
SW: sea water
PBST: phosphate buffered saline + 0.01% Triton X
NGS: normal goat serum
MOPS: 3-(N-Morpholino)propanesulfonic acid sodium salt
DMSO: Dimethyl sulfoxide
BSA: bovine serum albumin
DAPI: 4′,6-diamidino-2-phenylindole
DABCO: 1,4-diazabicyclo[2.2.2]octane)

## Declarations

### Ethics approval and consent to participate

not applicable

### Consent for publication

all authors approved the final manuscript for publication

### Availability of data and material

the datasets used during the current study are available from the corresponding authors on reasonable request.

## Authors’ contribution

PR, MS and GG: study design; MS, BB and SG: animal collection and embryo culture; RP, MS and GG: experiment design; MS, GG: experimental techniques optimization; PR and MS: phalloidin staining; MS: light microscopy; GG, MRE: transcriptome mining and phylogenetic analysis; MS, GG, PR: whole mount immunolocalization; MS, GG: *in situ* hybridization; MS, GG and AM: confocal acquisition; SG: SEM analyses; PR, MS, GG, AM, MRE and EBG: data interpretation; MS and GG: writing first draft of the manuscript; all authors: feedback on and approval of manuscript final version.

