## Supplementary material for "A feather star is born: embryonic development and nervous system organization in the crinoid *Antedon mediterranea*": MercurioGattoni_SupplementaryFigures

**Supplementary Figures:**

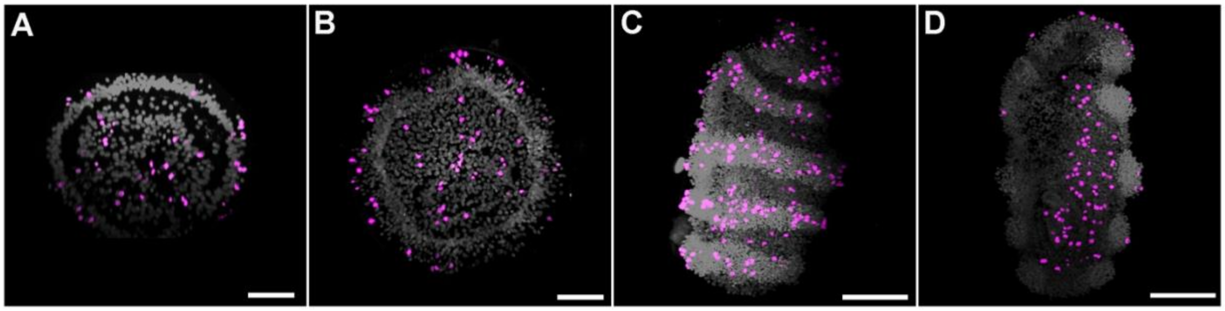


**Suppl. Fig. 1. Cell proliferation during post-cleavage stages in *A. mediterranea* embryos.** Confocal z-projections of embryos immunolabeled with PhH3 antibody (magenta) and DAPI (gray). Mid-sagittal sections of a representative mid-gastrula (20 hpf; **A**) and uniformly ciliated larva (48 hpf; **B**); lateral (**C**) and sagittal (**D**) projections of a pre-hatching larva (72 hpf). Scale bars: A and B = 50 µm; C and D = 100 µm.

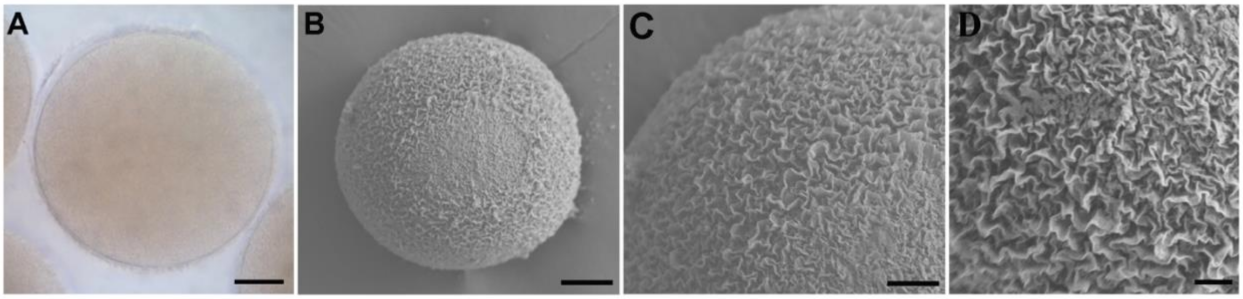


**Suppl. Fig. 2. Zygote of *A. mediterranea*. A**) Light microscopy of a zygote (~ 1 hpf) surrounded by its fertilization membrane; **B**) scanning electron microscopy of a zygote showing the irregular appearance of the fertilization membrane; **C** and **D**) magnification of B in which the thin ridges distributed along the surface of the embryo are observable. Scale bars: A = 50 µm; B = 50 µm; C = 20 µm; D = 10 µm.

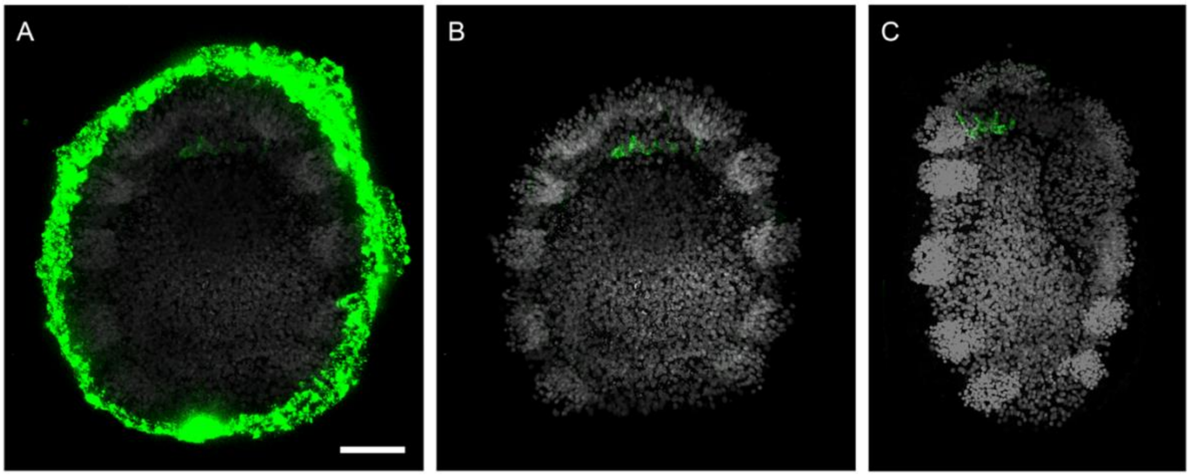


**Suppl. Fig. 3. Development of serotonergic neurons in pre-hatching stage.** Confocal z-projections showing serotonin immunoreactivity (green) and DAPI staining (grey) at the pre-hatching stage. Strong non-specific signal is found in the chorion membrane that surrounds the embryo before hatching (**A**). This signal was removed during image processing to reveal serotonergic cells in the developing apical organ (**B**, longitudinal section; **C**, sagittal section). Scale bar: 50µm

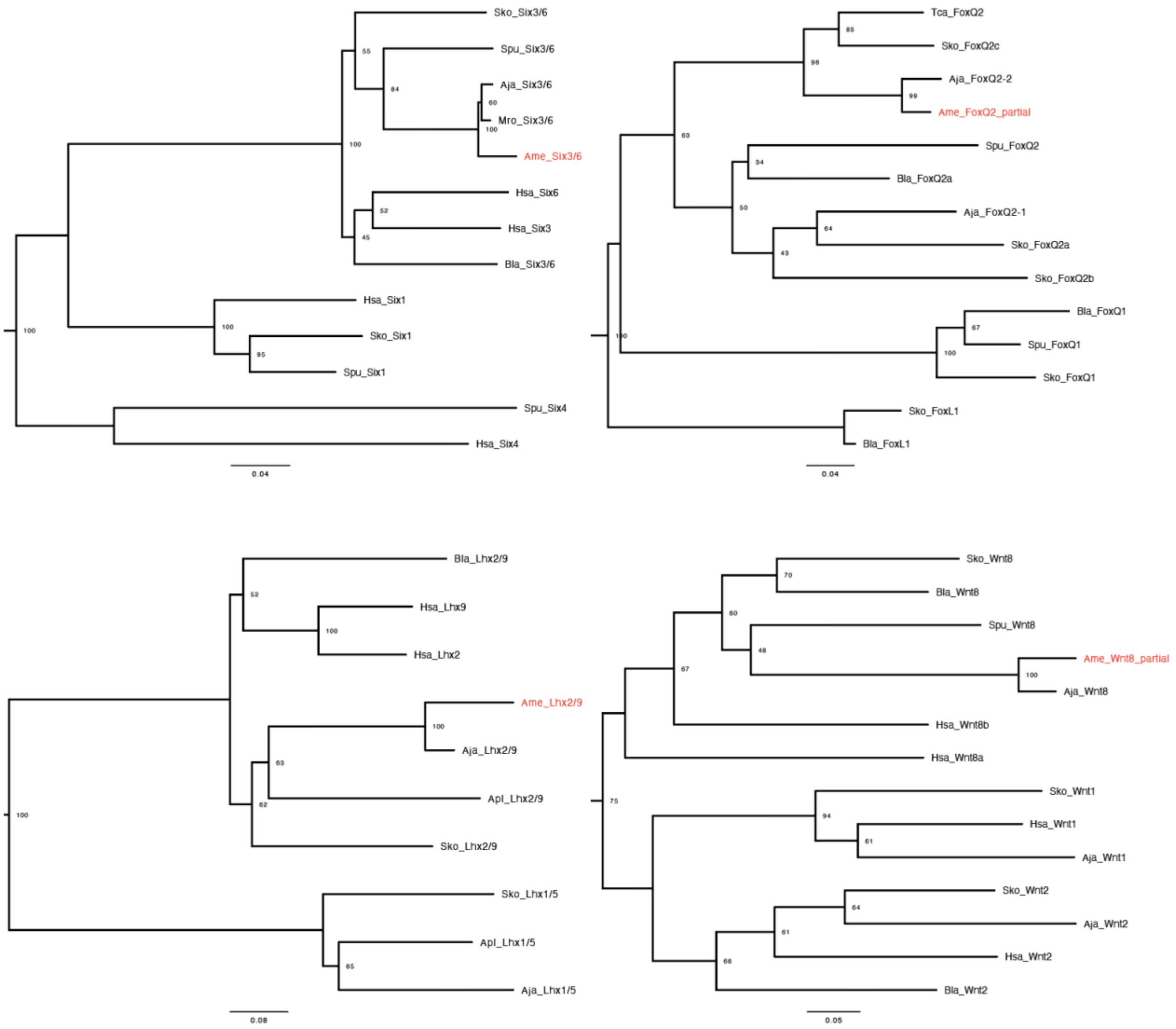


**Suppl. Fig. 4. Phylogenetic analysis of the relationships of *A. mediterranea FoxQ2, Six3/6, Lhx2/9* and *Wnt8* with related proteins in other taxa.** Phylogenetic trees built using the Bio-Neighbour-Joining method to determine the relationships of sequences recovered from the *A. mediterranea* transcriptome (shown in red) with related proteins in other taxa. Abbreviations: Aja: *Anneissia japonica*; Ame: *Antedon mediterranea*; Apl: *Acanthaster planci*; Bla: *Branchiostoma lanceolatum*; Has: *Homo sapiens*; Mro: *Metacrinus rotundus*; Sko: *Saccoglossus kowalevskii*; Spu: *Strongylocentrotus purpuratus* Tca: *Tribolium castaneum*;

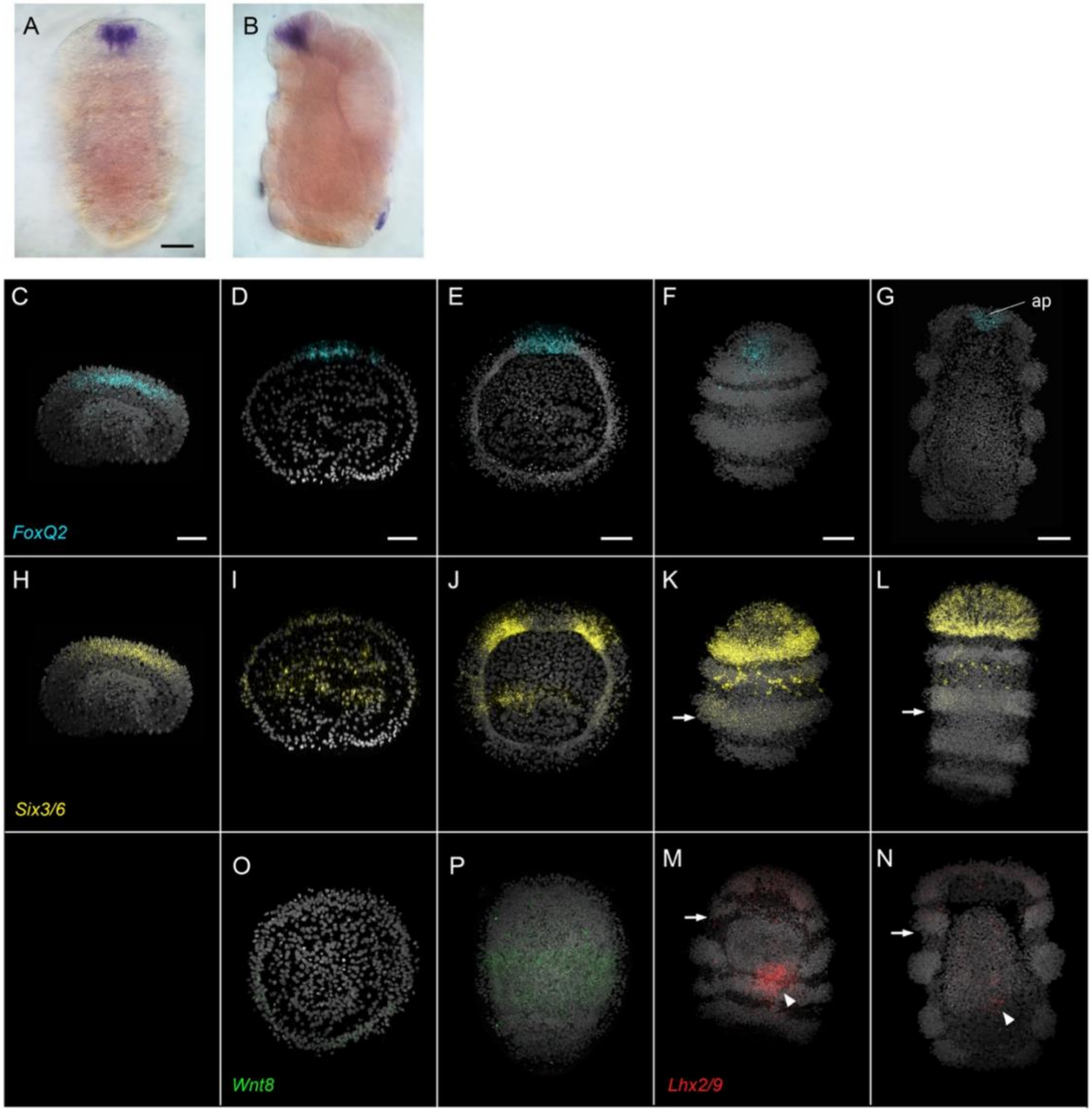


**Suppl. Fig. 5. Developmental expression of *Ame_FoxQ2*, *Ame_Six3/6*, *Ame_Lhx2/9* and *Ame_Wnt8* during *A. mediterranea* development.** (**A-B**) Expression of *Ame_FoxQ2* in the apical pit and apical organ of doliolaria larvae detected with chromogenic *in situ hybridization*. (**C-P**) Confocal z-projections showing expression of *Ame_FoxQ2* (cyan, **C-G**), *Ame_Six3/6* (yellow, **H-L**), *Ame_Lhx2/9* (red, **M-N**) and *Ame_Wnt8* (green, **O-P**) visualized through *in situ* hybridization chain reaction. Five developmental stages are shown: early gastrula (**C**, **H**), late gastrula (**D**, **I**, **O**), uniformly ciliated (**E**, **J**, **P**), pre hatching (**F**, **K**, **M**) and doliolaria (**G**, **L**, **N**) stages. Scale bar: 50μm

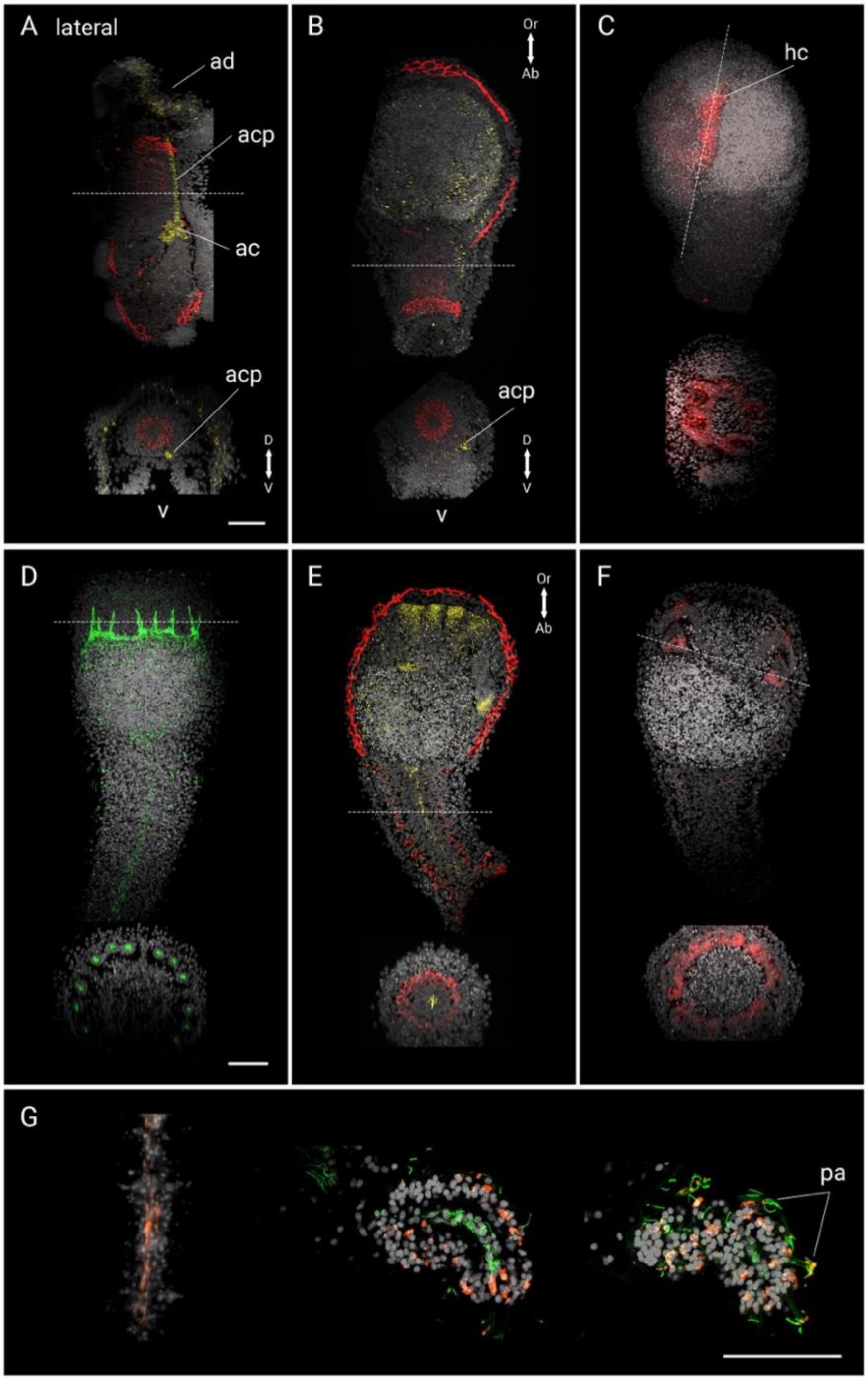


**Suppl. Fig. 6. Morphological and molecular characterization of crinoid metamorphosis.** Confocal projections showing localization of PSMAD1/5/8 (red) (**A**, **B**, **E**), acetylated α-tubulin (**D, G**) and glutamate (**G**) immunoreactivity and expression of *Ame_Six3/6* (yellow) (**A**, **B**, **E**) and *Ame_Lhx2/9* (red) (**C**, **F**) in doliolaria (**A**), settled (**B**-**C**), cystidean (**D**-**F**) and pentacrinoid (**G**)

stages. **A**) In the doliolaria stage, *Ame_Six3/6* marks the apical portion of the ectoderm as well as the axocoel and its anterior projection, which is located outside of the future stalk ossicles in the mesenchyme. **B**) After settlement, the ectodermal expression is lost while *Ame_Six3/6* remains expressed in the axocoel and its projection. **C**) At this stage, *Ame_Lhx2/9* is also lost from the ectoderm but its expression remains in the hydrocoel, which already shows a pentaradial organization. **D**) During metamorphosis, most of the organs undergo a 90° rotation, so that the vestibule and the hydrocoel are localized one below the other on the oral surface of the cystidean stage. Then, five sets of three projections direct orally from the hydrocoel and "push" on the vestibular ectoderm, forming the first 15 tube feet. As the hydrocoel is highly ciliated, these projections are visible with acetylated α-tubulin staining. Moreover, the somatocoels are also ciliated and can be seen surrounding the enteric sac and elongating into the stalk in the centre, surrounded by the developing stalk nerve. **E**) Interestingly, at this stage *Ame_Six3/6* is also expressed in the centre of the stalk, indicating that it starts to be expressed in the right somatocoel. This domain is different from the previous axocoelic one, as it is found within the ossicles. **F**) On the medial (ie closer to the centre) wall of the hydrocoel, at the base but not the tip of the forming tube feet, strong *Ame_Lhx2/9* signal could be detected. Both *Ame_Six3/6* and *Ame_Lhx2/9* started to be highly expressed on the oral surface of the vestibule, at the level of the external surface of the tube feet (**E-F**). **G**) Additional details of glutamatergic neurons distribution in the pentacrinoid. In the tube feet, glutamatergic cells are found in the ectoderm that surrounds the ciliated hydrocoel, and in the papillae from which short cilia emerge. Moreover, sparse glutamatergic cells are found in the stalk nerve. Legend: ac = axocoel: acp = axocoel projection; adp = adhesive pit; hc = hydrocoel; pa = papillae; rsc: right somatocoel; sn = stalk nerve; tfh: tube feet hydrocoel. Scale bar: 50µm
